# Accurate inference in parametric models reshapes neuroscientific interpretation and improves data-driven discovery

**DOI:** 10.1101/2020.04.10.036244

**Authors:** Pratik S. Sachdeva, Jesse A. Livezey, Maximilian E. Dougherty, Bon-Mi Gu, Joshua D. Berke, Kristofer E. Bouchard

## Abstract

A central goal of systems neuroscience is to understand the relationships amongst constituent units in neural populations and their modulation by external factors using high-dimensional and stochastic neural recordings. Statistical models, particularly parametric models, play an instrumental role in accomplishing this goal, because their fitted parameters can provide insight into the underlying biological processes that generated the data. However, extracting conclusions from a parametric model requires that it is fit using an inference procedure capable of selecting the correct parameters and properly estimating their values. Traditional approaches to parameter inference have been shown to suffer from failures in both selection and estimation. Recent development of algorithms that ameliorate these deficiencies raises the question of whether past work relying on such inference procedures have produced inaccurate systems neuroscience models, thereby impairing their interpretation. Here, we used the Union of Intersections, a statistical inference framework capable of state-of-the-art selection and estimation performance, to fit functional coupling, encoding, and decoding models across a battery of neural datasets. We found that, compared to baseline procedures, UoI inferred models with increased sparsity, improved stability, and qualitatively different parameter distributions, while maintaining predictive performance across recording modality, brain region, and task. Specifically, we obtained highly sparse functional coupling networks with substantially different community structure, more parsimonious encoding models, and decoding models that rely on fewer single-units. Together, these results demonstrate that accurate parameter inference reshapes interpretation in diverse neuroscience contexts. The ubiquity of model-based data-driven discovery in biology suggests that analogous results would be seen in other fields.

## 1 Introduction

Neuroscience, like many fields of biology, is undergoing a rapid growth in the size and complexity of experimental and observational data [1, 2]. Realizing the benefits of these advances in data acquisition requires improvements in the statistical models characterizing the data, as well as the inference procedures used to fit those models [3, 4]. For example, parametric models, such as generalized linear models, are appealing because the model parameters can be interpreted to gain insight into the underlying biological processes that generated the data [5–8]. However, even for this ubiquitously used class of models, the impact of an inference procedure’s statistical properties on biological interpretation is poorly appreciated.

These issues are particularly salient in systems neuroscience, where parametric models are often used to understand how neural activity is modulated by external factors (e.g., stimuli or a behavioral task) and internal factors (e.g., other neurons) [9, 10]. The fitted parameter values, therefore, specify which factors are important in modulating neural activity, and how important they are. The specific relationships that a parametric model describes ultimately frames how the model will be interpreted in a neuroscientific context, emphasizing the importance of accurate parameter inference.

For example, functional coupling models (Fig 1a) capture the statistical dependencies between different functional units in the brain, at scales ranging from single units to functional areas [5–8, 11–14]. These models can be used to construct networks [15, 16], which could be analyzed with an assortment of tools from graph theory to characterize the population [17, 18], related to structural connectivity [19], used to assess directed influence amongst neurons (i.e., effective connectivity) [20, 21], or related to external factors such as behavior, genetics, aging, or psychiatric conditions [22–24]. Encoding models map the dependence of a brain signal (e.g., neuronal spikes) on external factors, such as stimuli (Fig 1b) [25–27]. An example encoding model is a spatio-temporal receptive field of, for example, a visual cortex neuron, which maps the image space to the neuronal response (Fig 1b, right) [28–30]. More refined encoding models of neural population data can be used to test theoretical and computational theories of neural coding [31–33]. On the other hand, decoding models map the brain signal to external factors, using the activities in, e.g., a neural population, to predict a stimulus or task-relevant behavioral condition (Fig 1c) [34–40]. A common linear decoding model is the extraction of a hyperplane in the neural activity space, which provides a decision boundary for one of two behavioral conditions or stimuli (Fig 1c, right: *s*_1_, *s*_2_) [22, 41, 42], though recent work has explored more advanced decoders, using artificial neural networks [39] or predictive latent representations [43]. Using decoding models for brain-computer interfaces has both clinical uses and scientific implications for understanding learning and motor control [44, 45]. Since these models are used to make scientific conclusions about the function of the brain, understanding the stability, accuracy, and parsimony of the inference procedures and resulting models is of paramount importance.

**Figure 1:**
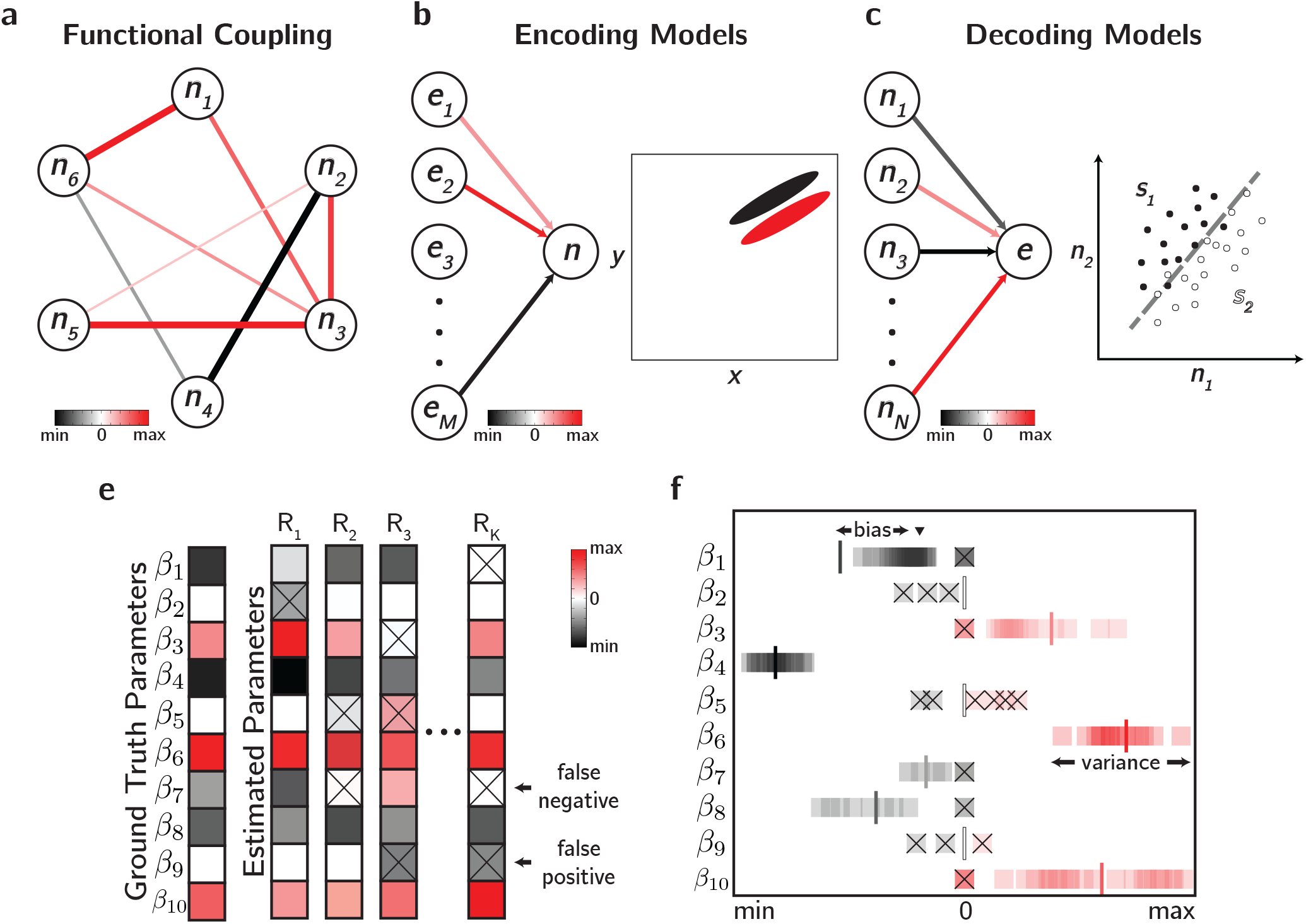
Parametric models and statistical inference in systems neuroscience. **a-c.** Three examples of parametric models widely used in systems neuroscience. **a.** Functional coupling models characterize the statistical relationships between neurons in a population. **b.** Encoding models map *M* internal or external factors 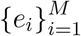 to a neuronal response *n*. **c.** Decoding models map the activities of *N* neurons in a population 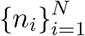 to an internal or external factor *e*. **e-f.** Quantification of statistical selection and estimation performance. **e.** The ground truth values of the parameters in an example model and estimated values across *K* different resamples of the data. **f.** The distribution of estimated values for each parameter in the ground truth model of the previous panel, with the true values denoted by vertical lines. For *β*_1_, ▾ denotes the mean estimated value across resamples. False positives and false negatives are denoted with an ×.

The utility of parametric models hinges on the assumption that the inference procedure used to fit them selects the correct parameters (i.e., specified as zero or non-zero) and properly estimates their values. The statistical consequences of improper selection are false positives or false negatives (Fig 1d), while poor estimation results in high bias (Fig 1e: e.g., *β*_1_) or high variance (Fig 1e: e.g., *β*_6_). The neuroscientific consequences of statistical inference lie in the interpretation of the fitted parametric model. Selection informs which internal and external factors are relevant for predicting neural activity, and estimation specifies their relative importance. Thus, validating that an inference procedure can reliably select and estimate a model’s parameters is vital to ensure that they motivate correct conclusions about neural activity.

These issues imply that, when fitting parametric models in a scientific context, multiple goals beyond predictive performance must be balanced to produce a scientifically meaningful model. In particular, achieving a parsimonious model, which uses the fewest number of features to sufficiently predict the response variable (i.e., finding the “simplest”), has long served as a goal in statistical model selection [46]. One approach to model parsimony relies on the imposition of sparsity during feature selection, which has the added benefit of identifying a small subset of predictive features, facilitating the interpretability of the model [47, 48]. This is particularly relevant in high-dimensional settings where there are few task-relevant features and strong priors from domain knowledge for selection may not exist. Another desired property is stability, or the reliability of a machine learning algorithm when its inputs are slightly perturbed [49, 50]. For a model to be interpretable, its parameters must be robust to the often noisy processes that generated the data. Thus, encouraging stability in a model’s parameter inference procedure will ensure that the features describing the relevant signal are selected and their correct contributions are properly estimated [51, 52]. Until recently, inference procedures that sufficiently balanced accurate selection and estimation, predictive performance, and stability were lacking.

The recently introduced Union of Intersections (UoI) is a statistical inference framework based on stability principles which enhances inference in a variety of common parametric models [53]. The properties characterizing UoI models — sparsity, stability, and predictive accuracy — are well-suited to data-driven discovery in neuroscience, due to the high dimensionality and many sources of variability in its datasets. Thus, UoI can be leveraged to assess whether baseline approaches to parameter inference in common models are susceptible to improper feature selection and estimation, and if so, assess the consequences for model interpretability in a neuroscience context.

In this work, we used the UoI framework to fit functional coupling, encoding, and decoding models to diverse neural data in an effort to elucidate the impacts of precise selection and estimation on neuroscientific interpretation. We found that, compared to baseline procedures, we obtained models with enhanced sparsity, improved stability, and significantly different parameter distributions, while maintaining predictive performance across recording modality, brain region, and task. Specifically, we obtained highly sparse coupling models of rat auditory cortex, macaque V1, and macaque M1 without loss in predictive performance. These models were used to construct functional networks that exhibited enhanced modularity and decreased small-worldness. We built parsimonious encoding models of mouse retinal ganglion cells and rat auditory cortex that more tightly matched with theory. These models were able to predict held-out neural responses with parameters that were as simple as possible, but no simpler. Lastly, we decoded task-relevant external factors from rat basal ganglia activity using fewer single units than baseline models. Overall, by emphasizing accurate selection and estimation during the inference of parametric neural models, we constructed equally predictive models that reshaped neuroscientific interpretation across a diverse set of neural data.

## 2 Methods

Our goal is to demonstrate how the statistical properties of inference algorithms impact the fitting and interpretation of diverse parametric models commonly used in neuroscience. The main tools we use for this purpose are algorithms based on the Union of Intersections framework [53–55]. Given the novelty of Union of Intersections, we first describe it conceptually, which also serves to introduce other relevant background. We then briefly summarize a comparison of a specific UoI algorithm, UoI_Lasso_, versus other algorithms on a synthetic dataset to motivate UoI as a superior selection and estimation algorithm for subsequent application to neural data in the Results section.

### 2.1 The Union of Intersections framework balances sparsity, stability, and predictive performance

Union of Intersections (UoI) is not a single method or algorithm, but a flexible framework into which other algorithms can be inserted for enhanced inference. In this work, we apply the UoI framework to generalized linear models, focusing on linear regression (UoI_Lasso_), Poisson regression (UoI_Poisson_) and logistic regression (UoI_Logistic_). We refer the reader to UoI variants of other procedures, such as non-negative matrix factorization [54] and column subset selection [53].

Consider the general problem of mapping a set of *p* features **x**∈ ℝ^*p×*1^ to a response variable *y* ∈ ℝ, of which we have *N* samples 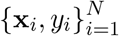. For convenience, we focus on linear models, which require estimating *p* parameters **β** ∈ ℝ^*p×*1^ that linearly map **x**_*i*_ to *y*_*i*_. We describe the UoI framework in this context, which involves the algorithm UoI_Lasso_. The steps we detail, however, extend naturally to other penalized generalized linear models [56]. Typically, the mapping in linear models is corrupted by i.i.d. Gaussian noise *ϵ*:

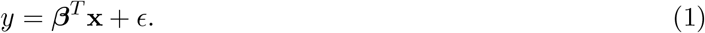

The parameters *β* can be inferred by optimizing the traditional least squares error on *y*:

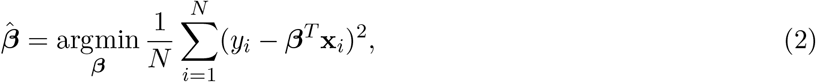

where *i* indexes the *N* data samples. The UoI framework combines two techniques — regularization and ensemble methods — to balance sparsity, stability, and predictive performance, thereby improving on the traditional least squares estimate (Fig 2a).

**Figure 2:**
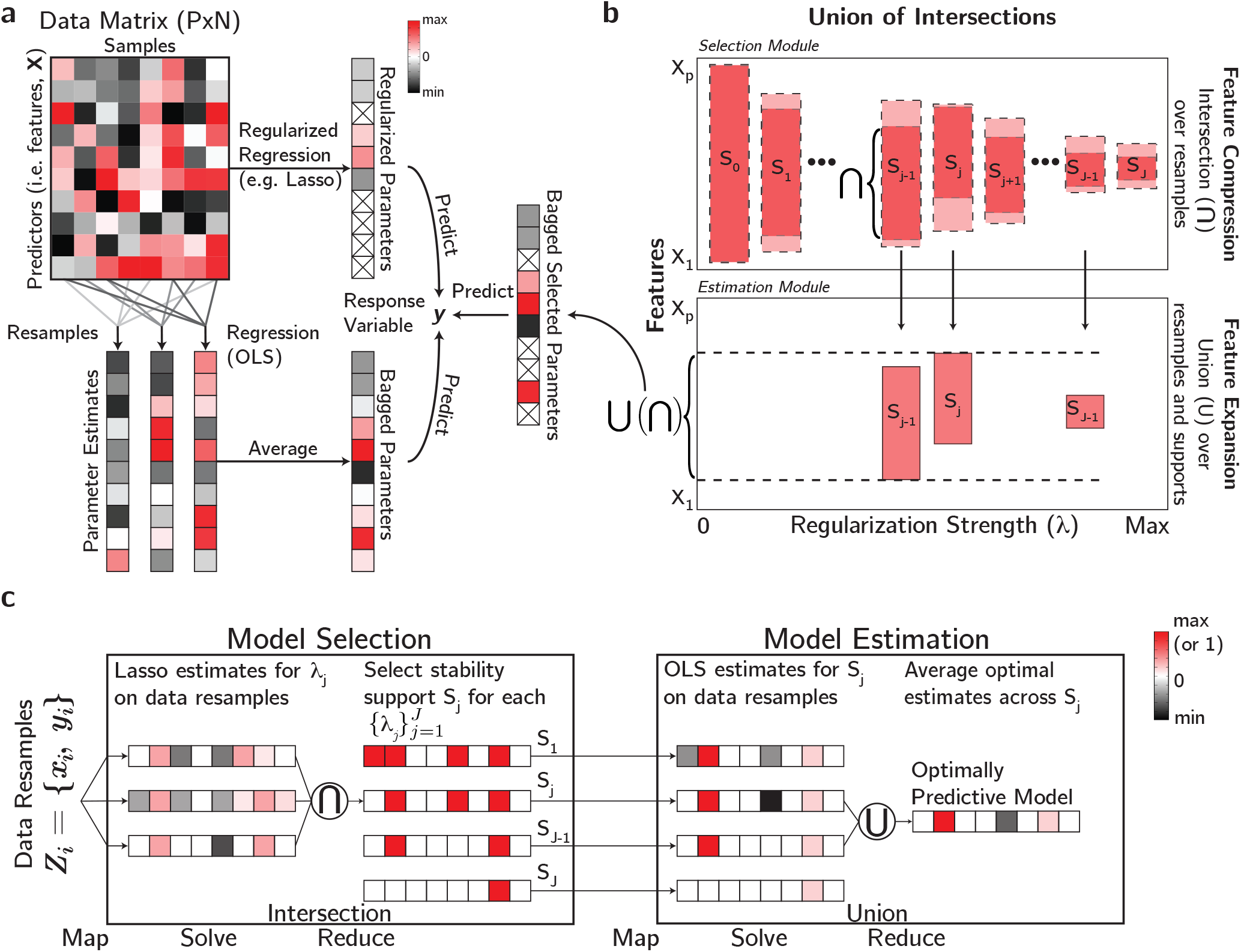
The Union of Intersections framework combines ensemble and regularization approaches in model inference. **a.** Schematic of regularization and ensemble methods. Top: Regularization can be used to perform feature selection. Features set exactly to zero are denoted by ×. Bottom: Ensemble procedures aggregate model fits across resamples of the data. **b.** Schematic of the UoI framework. The *x*-axis corresponds to the set of regularization parameters while the *y*-axis corresponds to the feature space (*X*_1_, *X*_2_, … , *X*_*p*_). Pink bands denote features included in a support or estimated model. Dark pink bands denote features included after the intersection step. **c.** UoI_Lasso_ depicted in a data-distributed fashion. *Model Selection (left):* Left column depicts the lasso estimates across data resamples for a specific hyperparameter *λ*_*j*_. Right column depicts the intersected supports for different *λ*_*j*_, with the second support referring specifically to the estimates in the left column. Red here denotes 1, rather than maximum value. *Model Estimation (right):* Left column depicts the parameter estimates for one data resample, per support. Models with the best predictive performance (second and third rows) are unionized to obtain the final model (right column).

Structured regularization, or the inclusion of penalty terms in the objective function to restrict the model complexity, can be useful when a subset of the *β*_*i*_ are exactly equal to zero, i.e., *β* is sparse. Sparsity implies that some features are not relevant for predicting the response variable. This assumption is often useful for data-driven discovery in biological settings, particularly for framing the interpretation of the model in the context of physical processes that generated the data. The identification of which *β*_*i*_ are non-zero can be viewed as a feature selection (or more generally, model selection) problem [48]. A common regularization penalty used for feature selection is the lasso penalty |*β*|_1_, or the *ℓ*_1_-norm applied to the parameters [47]. For the case of linear regression, this creates an optimization problem of the form

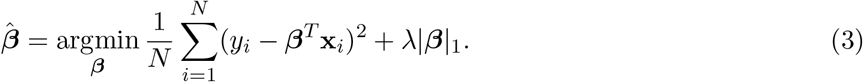

Solving Eq (3) returns parameter estimates with some sparsity, provided that *λ* is appropriately chosen (Fig 2a, top). Typically, *λ*, the degree to which feature sparsity is enforced, is unknown and must be determined through cross-validation or a penalized score function such as the Bayesian information criterion (BIC) [46] across a set of *J* hyperparameters 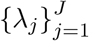. Importantly, solving the lasso problem simultaneously performs model selection (identifying the non-zero features) and model estimation (determining the specific values of those parameters). However, the application of the lasso penalty suffers from shrinkage [47], or a parameter bias that erroneously reduces the magnitudes of the parameters (Fig 2a, top: compare opacity of parameter estimates), and often does not correctly identify the true non-zero parameters (Fig 2a, top: false positives).

On the other hand, ensemble procedures (e.g., bagging and boosting [57, 58]) aggregate model fits across resamples of the data to improve the stability of parameter estimates (Fig 2a, bottom). The more stable parameter estimates result in improved predictive accuracy. This is particularly desirable in biological settings, where model aggregation ensures that the relevant signal in noisy data is reflected in the parameter estimates. However, ensemble procedures do not perform feature selection.

UoI separates model selection and model estimation into two stages, with each stage utilizing ensemble procedures to promote stability. Specifically, model selection is performed through intersection (compressive) operations and model estimation through union (expansive) operations, in that order. This separation of parameter selection and estimation provides selection profiles that are robust and parameter estimates that have low bias and variance. Fig 2b and 2c provide a visual depiction of the UoI framework. For UoI_Lasso_, the procedure is as follows:

#### Model Selection

Define the support *S* as the set of non-zero parameters in an estimate 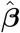. For each λ_*j*_, generate parameter estimates by solving the lasso optimization problem (Eq 3) on *N*_*S*_ resamples of the data, and calculate a support for each resample-λ_*j*_ pairing. The intersection step requires that only the features that appear in a sufficient number of resamples are included in the final *stability support S*_*j*_ for λ_*j*_. We depict this in Fig 2b, top, where the light pink bands denote features that are included in the model due to regularization while the dark pink bands denote features that are included in the stability support after the intersection across resamples. The bands are arranged in order of increasing regularization strength (Fig 2b: *x*-axis) and thus sparser (i.e., smaller) support sets (Fig 2b: *y*-axis). Note that the stability support may be calculated with a hard intersection (e.g., Fig 2c: Model Selection) or a soft intersection. In the former case, a feature must appear in the support of every resample to be included in *S*_*j*_. In the latter, the feature must only appear in a sufficient fraction of supports which is a hyperparameter.

#### Model Estimation

For each stability support *S*_*j*_, estimate the model without regularization on *N*_*E*_ resamples of the data. Each resample contributes the (now fitted) support that had the best predictive performance on its data to the union step (Fig 2b: bottom). Unique supports may have the best performance across multiple resamples (e.g., only three unique supports, *S*_*j*−1_, *S*_*j*_, and *S*_*m*−1_, are included for model averaging in Fig 2b). The best fitted parameter estimates for each resample are unionized according to some metric (e.g., median, mean, etc.), resulting in a final parameter estimate (Fig 2b, bottom). For UoI_Lasso_, this procedure consists of applying Ordinary Least Squares to each stability support and resample combination (Fig 2c: Model Estimation).

UoI’s modular approach to parameter inference capitalizes on the feature selection achieved by stability selection and the unbiased, low-variance properties of the bagged OLS estimator. Furthermore, UoI’s novel use of model aggregating procedures within its resampling framework allows it to achieve highly sparse (i.e., only using features robust to perturbations in the data) and predictive (i.e., only using features that are informative) model fitting. Importantly, this is achieved without imposing an explicit prior on the model distribution, and without formulating a non-convex optimization problem. Since the optimization procedures across resamples can be performed in parallel, the UoI framework is naturally scalable, a fact that we have leveraged to facilitate parameter inference on larger datasets [55]. The application of UoI_Lasso_ in a data-distributed manner is depicted in Fig 2c. In the selection module, the first column depicts lasso estimates across data resamples for a particular choice of regularization parameter, all of which can be fit in parallel (Fig 2c, Model Selection: left column). In the estimation module, OLS estimates are fit across resamples and supports, which can be done in parallel (Fig 2c, Model Estimation: left column).

### 2.2 Model evaluation

We describe the metrics used to evaluate the models fitted to synthetic data and the neural data. Note that only the selection ratio, predictive performance measures, and Bayesian information criterion were used to evaluate models on neural data where ground truth was unknown.

#### Selection accuracy

The selection accuracy, or set overlap, is a measure of how well the estimated support captures the ground truth support. Define *S*_*β*_ as the set of features in the ground truth support, 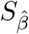 as the set of features in the estimated model’s support, |*S*| as the cardinality of *S*, and Δ as the symmetric set difference operator. Then the selection accuracy is defined as

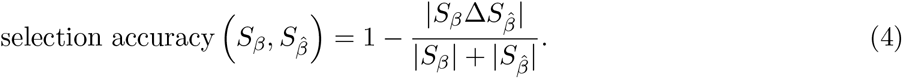

The selection accuracy is bounded in [0, 1], taking value 0 if 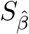 and *S*_*β*_ have no elements in common, and taking value 1 iff they are identical.

#### Estimation error

The estimation error of the *p* fitted parameters 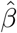, with ground truth parameters *β*, is defined as the root mean square error, or

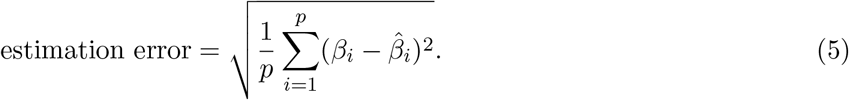

#### Estimation variability

The estimation variability for parameter *β*_*i*_ is defined as the parameter standard deviation *σ*(*β*_*i*_). We calculated this quantity by taking the variance of the estimated parameter 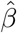 over *R* resamples of the data:

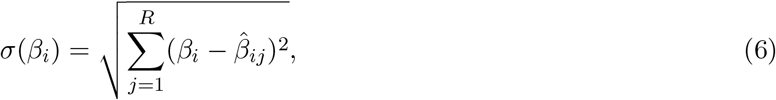

where *j* indexes the resample. To summarize this measure across all *p* parameters in a model, we took the average, i.e., 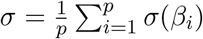.

#### Selection ratio

We evaluate the sparsity of estimated models with the selection ratio, or the fraction of parameters fitted to be non-zero:

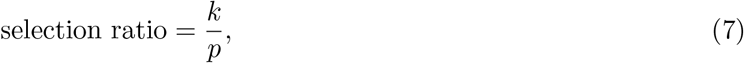

where *p* is the total number of parameters available to the model and *k* is the number parameters that a model-fitting procedure explicitly sets non-zero.

#### Predictive performance

We utilized several measures of predictive performance, depending on the model. For linear models (i.e., a generalized linear model with an identity link function), we used the coefficient of determination (*R*^2^) evaluated on held-out data:

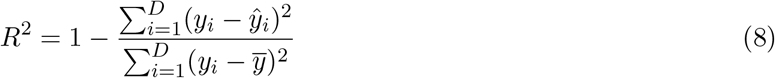

where *y*_*i*_ is the ground truth response for sample *i*, 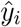 its corresponding predicted value, and 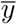 the mean of the response variable over trials. *R*^2^ has a maximum value of 1, when the model perfectly predicts the response variable across samples. *R*^2^ values below zero indicate that the model is worse than an intercept model (i.e., simply using the mean value to predict across samples).

For Poisson regression, or a generalized linear model with a logarithmic link function, we utilized the deviance, which is the difference in log-likelihood between the saturated model and the estimated model [59]. The saturated model has parameters specifically chosen to reproduce the observed values. For the Poisson log-likelihood, the expression for the deviance as a function of the estimation parameters 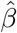 is given by

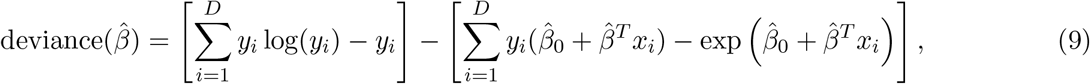

where 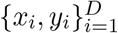 denote the features and response variable of the model, respectively. Note that lower deviance is preferred, in contrast to the coefficient of determination.

For logistic decoding models, we used the classification accuracy on held-out data as the measure of predictive performance.

#### Model Parsimony

We evaluated model parsimony using the Bayesian information criterion (BIC) [46]:

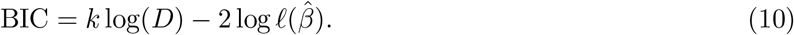

Here, *D* is the number of samples, *k* is the number of parameters estimated by the model, and log 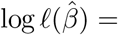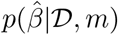 is the log-likelihood of the parameters 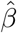 under data 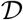 and model *m*. Thus, the BIC includes a penalty that encourages models to be more sparse (first addend) while still accounting for predictive accuracy (second addend). Importantly, the BIC is evaluated on the data that the model was trained on (rather than held-out data). It is typically used as a model selection criterion (in lieu of, for example, cross-validation). When used as a model selection criterion, the model with lower BIC is preferred.

### 2.3 UoI_Lasso_ achieves superior selection and estimation performance on synthetic data

We evaluated UoI_Lasso_’s abilities as an inference procedure by assessing its performance on synthetic data generated from a linear model. The performances of UoI and five other inference procedures are depicted in Fig 3: UoI_Lasso_ (black), ridge regression (purple) [48], lasso (green) [47], smoothly clipped absolute deviation (SCAD; red) [60], bootstrapped adaptive threshold (BoATS; blue) [61], and debiased lasso (dbLasso; coral) [62].

**Figure 3:**
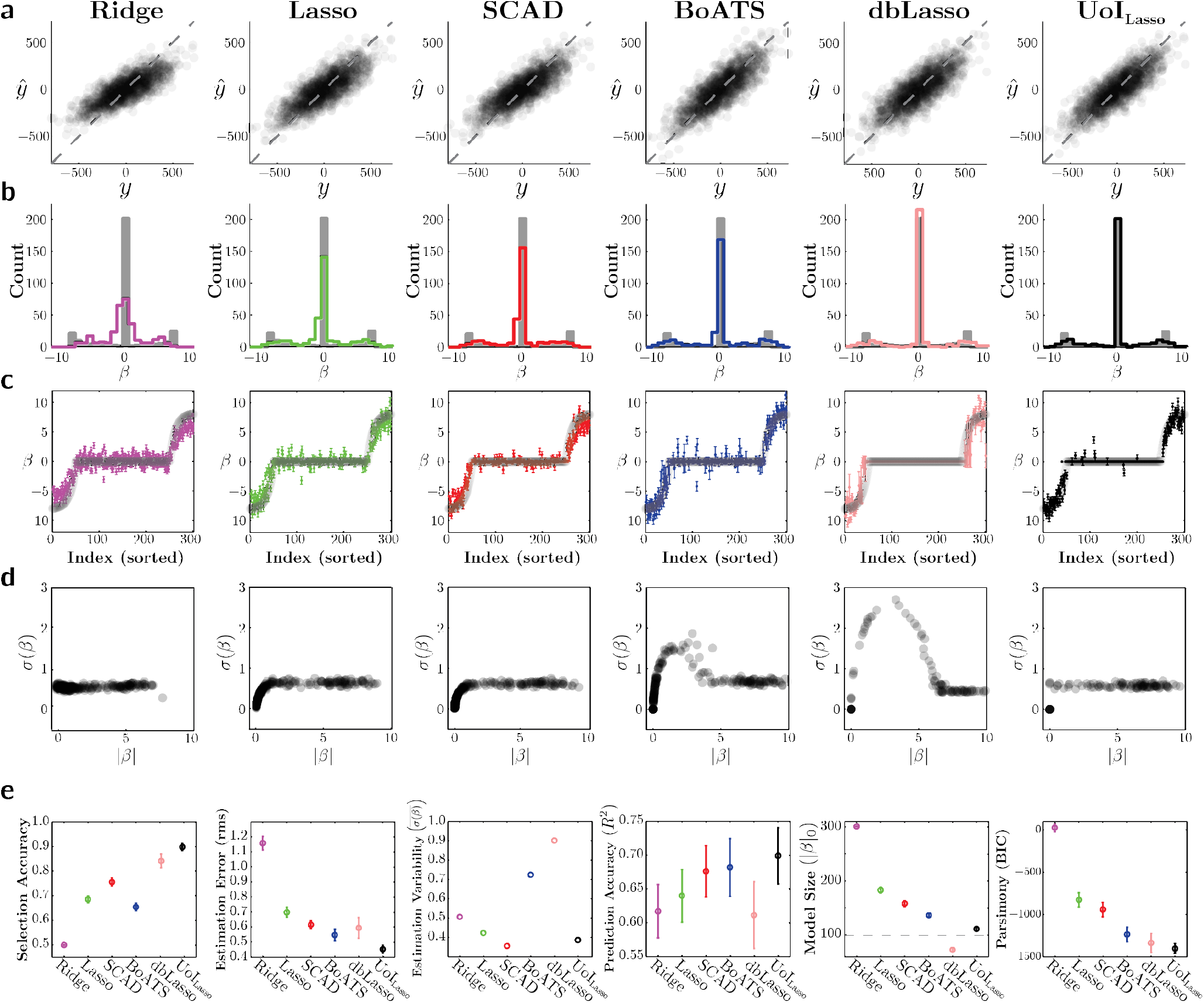
UoI achieves superior selection and estimation performance on synthetic data over a battery of alternative inference algorithms. The performance of ridge regression (purple), lasso regression (green), smoothly clipped absolute deviation (SCAD; red), bootstrapped adaptive threshold (BoATS; blue), debiased lasso (dbLasso; coral), and UoI_Lasso_ (black) on data generated from a synthetic linear model. Panels **a-d** highlight separate measures per row, with each column referring to a separate inference algorithm. Each column in panel **e.** directly compares a summary measure across inference algorithms. **a.** Comparison of the predicted and true values of the response variable for held-out data. **b.** Histogram of the estimated parameters (colored outline) compared to the distribution of true parameters (gray). **c.** Estimated parameters (colored points; error bars denote the IQR across data resamples) compared to true parameters (gray), both sorted on the *x*-axis according to the value of the true parameter. **d.** Variance of the parameter estimates across data resamples as a function of the mean estimated parameter’s magnitude. **e.** Direct comparison of the selection accuracy, estimation error (root mean square of parameter estimates), estimation variability (mean parameter variance), prediction accuracy (coefficient of determination), the inferred model’s size (number of identified non-zero parameters, with the black dashed line denoting ground truth), and parsimony (Bayesian information criterion). Error bars denote IQR across data resamples.

The linear model consisted of *p* = 300 total parameters, with *k* = 100 non-zero parameters (thereby having sparsity 1 − *k/p* = 2/3). The non-zero ground truth parameters were drawn from a parameter distribution characterized by exponentially increasing density as a function of parameter magnitude (Fig 3b: gray histograms). We used *N* = 1200 samples generated according to the linear model (1) with noise magnitude chosen such that Var(*ϵ*) = 0.2 × |*β*|_1_. We report metrics according to their statistics across 100 randomized cross-validation samples of the data.

In Fig 3a, we show scatter plots comparing the predicted and actual values of the observation variable on held-out data samples. We visualized how well the inference procedures captured the underlying parameter distribution by comparing the histograms of (average) estimated model parameters (colors) overlaid on the ground truth model parameters (grey) (Fig 3b). We additionally plotted parameter bias and variance, first by comparing the mean estimated value (± standard deviation) against the ground truth parameter value (Fig 3c), and then examining the standard deviation of the parameter estimates as a function of their mean estimated value (Fig 3d).

Fig 3a-d captures the improvements that the UoI framework offers in parameter inference. UoI_Lasso_ is designed to maximize prediction accuracy (Fig 3a) by first selecting the correct features (Fig 3b), and then estimating their values with high accuracy (Fig 3c) and low variance (Fig 3d). By separating model selection and model estimation, UoI_Lasso_ benefits from strong selection (as in BoATS and debiased Lasso), but with the low variability of the structured regularizers (Lasso, SCAD), while alleviating shrinkage with its nearly unbiased estimates.

We quantified the performance of the inference algorithms on synthetic data using a variety of metrics capturing selection performance, bias, variance, and prediction accuracy. UoI_Lasso_ generally resulted in the highest selection accuracy (Fig 3e, first column), parameter estimates with lowest error (Fig 3e, second column) and competitive variance (Fig 3e, third column). In addition, it led to the best prediction accuracy (Fig 3e, third column). UoI_Lasso_ best captured the true model size (Fig 3e, fourth column), avoiding the abundance of false positives suffered by most other inference algorithms. UoI_Lasso_’s enhanced predictive performance with fewer features resulted in superior model parsimony (Fig 3e, fifth column).

### 2.4 Neural recordings

We sought to demonstrate impact of improved inference on parametric models across a diversity of datasets, spanning distinct brain regions, animal models, and recording modalities. We used microelectrocorticography recordings obtained from rat auditory cortex (for coupling and encoding models), single-unit recordings from macaque visual and motor cortices (coupling models), single-unit recordings from isolated rat retina (encoding models), and single-unit recordings from basal ganglia (decoding models). We briefly describe the experimental and preprocessing steps for each dataset.

#### Recordings from auditory cortex

Auditory cortex (AC) data was comprised of cortical surface electrical potentials (CSEPs) recorded from rat auditory cortex with a custom fabricated micro-electrocorticography (*μ*ECoG) array. The *μ*ECoG array consisted of an 8 × 16 grid of 40 *μ*m diameter electrodes. Anesthetized rats were presented with 50 ms tone pips of varying amplitude (8 different levels of attenuation, from 0 dB to −70 db) and frequency (30 frequencies equally spaced on a log-scale from 500 Hz to 32 kHz). Each frequency-amplitude combination was presented 20 times, for a total of 4200 samples. The response for each trial was calculated as the *z*-scored, to baseline, high-*γ* band analytic amplitude of the CSEP, calculated using a constant-Q wavelet transform. Of the 128 electrodes, we used 125, excluding 3 due to faulty channels. Data was recorded by Dougherty & Bouchard (DB). Further details on the surgical, experimental, and preprocessing steps can be found in [63].

#### Recordings from primary visual cortex

We analyzed three primary visual cortex (V1) datasets, comprised of spike-sorted units simultaneously recorded in three anesthetized macaque monkeys. Recordings were obtained with a 10 × 10 grid of silicon microelectrodes spaced 400 *μ*m apart and covering an area of 12.96 mm^2^. Monkeys were presented with grayscale sinusoidal drifting gratings, each for 1.28 s. Twelve unique drifting angles (spanning 0° to 330°) were each presented 200 times, for a total of 2400 trials per monkey. Spike counts were obtained in a 400 ms bin after stimulus onset. We obtained [106, 88, 112] units from each monkey. The data was obtained from the Collaborative Research in Computational Neuroscience (CRCNS) data sharing website [64] and was recorded by Kohn and Smith (KS) [65]. Further details on the surgical, experimental, and preprocessing steps can be found in [66] and [67].

#### Recordings from primary motor cortex

Primary motor cortex (M1) data was comprised of spike-sorted units simultaneously recorded in the motor cortex of Rhesus macaque monkey. Recordings were obtained with a chronically implanted silicon microelectrode array consisting of 96 electrodes spaced at 400 *μ*m and covering an area of 16 mm^2^. We used three datasets, consisting of three recording sessions from monkey I. The behavioral task required the monkey to make self-paced reaches to targets arranged on a 8 × 17 grid. Spike counts were binned at 150 ms over the course of the entire recording session, resulting in [4089, 4767, 4400] samples per recording session. We obtained [136; 146; 147] units from each dataset. Data was recorded by O’Doherty et al. (OCMS) and obtained from Zenodo [68]. Further details on the surgical, experimental, and preprocessing steps can be found in [69].

#### Recordings from retina

Retina data comprised spiking activity, extracellularly recorded from isolated mice retina. Recordings were obtained using a 61-electrode array. Isolated retina were presented with a flicking black or white bar stimulus according to a pseudo-random binary sequence for a period of 16.6 ms. We utilized recordings from 23 different retinal ganglion cells. Data was obtained from CRCNS and recorded by Zhang et al [70]. Further details on the surgical, experimental, and preprocessing steps can be found in [71].

#### Recordings from basal ganglia

Basal ganglia data comprised tetrode recordings from two regions of rat basal ganglia: the globus pallidus pars externa (GPe: 18 units) and substantia pars nigra reticulata (SNr: 36 units). Recordings were performed during a rodent stop-signal task. Briefly, a rat was prompted to enter a center port with a light cue. The rat remained in the port until a Go cue (audio stimulus at 1 kHz or 4 kHz) which directed a lateral head movement to the left or right ports. On a subset of trials, the Go cue was followed by a Stop signal (white noise burst), indicating that the rat should remain in the center port. We utilized the successful Go trials, in which the rat was not given a Stop signal and successfully entered the correct port (186 trials). We used the spike count in the first 100 ms after the rat exited the center port to predict the behavioral condition (left or right). Further details on the surgical, experimental, and preprocessing steps can be found in [72].

### 2.5 Neural data analysis and model fitting

All models fit to neural data consisted of various generalized linear models, depending on the application. We trained all baseline models using either the `glmnet` [56] or `scikit-learn` [73, 74] packages. Meanwhile, we trained all UoI models using the `pyuoi` package [55].

#### Model selection criterion in the estimation module

In the UoI framework, the estimation module operates by unionizing fitted stability supports across resamples. Thus, the module requires a criterion by which to choose the best fitted stability support per resample. A natural choice, akin to cross-validation, is the out-of-resample validation performance according to some measure (e.g., *R*^2^, deviance, etc.). However, the use of cross-validation as a model selection tool is known to suffer from inconsistent feature selection [75–77]. Therefore, we instead utilized the Bayesian information criterion in the estimation module for each model, which has been shown to be model selection consistent [46].

#### Data analysis for coupling models

We used UoI_Lasso_ (rat auditory cortex) and UoI_Poisson_ (macaque V1 and M1) to fit coupling models. The auditory cortex model can be described with a linear model as

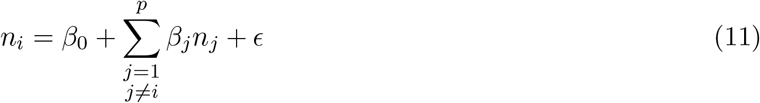

where *n*_*i*_ is the high-gamma activity of the *i*th electrode on a trial. The baseline procedure consisted of a lasso optimization with coordinate descent, while the UoI approach utilized UoI_Lasso_. The model for the spiking datasets, which utilizes a Poisson generalized linear model, can be written as

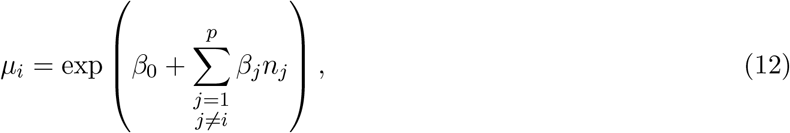

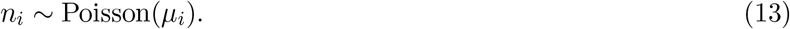

where *n*_*i*_ corresponds to the spike count of the *i*th neuron. The corresponding objective function for this model is the log-likelihood,

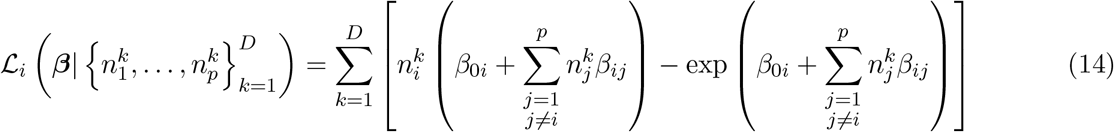

where *i* denotes that this model corresponds to the *i*th neuron, *j* indexes over the remaining neurons, and *k* indexes over the *D* data samples. The baseline approach consisted of applying coordinate descent to solve this objective function with an elastic net penalty:

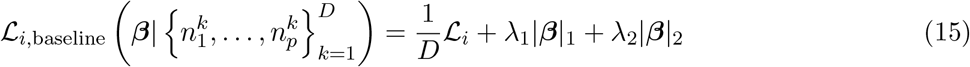

where λ_1_ and λ_2_ are the two hyperparameters specifying the strength of the *ℓ*_1_ and *ℓ*_2_ penalties, respectively. Note that the intercept terms were not penalized. Meanwhile, UoI_Poisson_ utilized objective function (14) with only an *ℓ*_1_ penalty in the selection module. In the estimation module, we used Eq (14) with a very small *ℓ*_2_ penalty for numerical stability purposes. The specific optimization algorithm was a modified orthant-wise L-BFGS solver [78].

#### Data analysis for encoding models

For retinal data, we fit spatio-temporal receptive fields (STRFs) frame-by-frame. Specifically, the STRF was comprised of *F* frames *β*_1_, *β*_2_, … , *β*_*F*_, each a vector of size *M* and spanning Δ*t* seconds. For neuron *i* and frame *k*, the encoding model consisted of

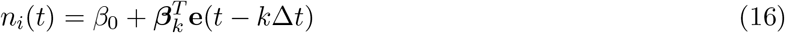

where *n*_*i*_(*t*) is the spike count at timepoint *t* and **e**(*t* − *k*Δ*t*) is flicking bar stimulus value at *k* bins before *t*. We fit the *F* models using lasso (baseline) and UoI_Lasso_, and created the final STRF by concatenating the parameter values *β* = [*β*_1_, **β**_2_, … , *β*_*F*_].

The tuning model for the rat auditory recordings was constructed using Gaussian basis functions. We used eight Gaussian basis functions spanning the log-frequency axis with means 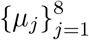 [5, 13]. Thus, the high-gamma activity *n*_*i*_ of electrode *i* in response to frequency *f* was

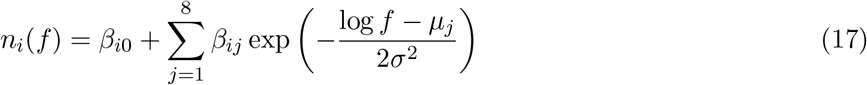

We chose *σ*^2^ = 0.64 octaves so that basis functions sufficiently spanned the plane. We chose *p* = 8 basis functions because this was the minimum number of basis functions for which every electrode had a selection ratio less than 1. We fit Eq (17) using cross-validated lasso as the baseline. To characterize the relationship between selection ratio and predictive performance of the rat AC tuning models, we fit trendlines across models using Gaussian process regression. Specifically, we utilized a regressor with radial basis function kernel (length scale *ℓ* = 0.01) and a white noise kernel (noise level *α* = 0.1).

#### Data analysis for decoding models

We fit decoding models to basal ganglia recordings as binary logistic regression models. The model expresses the probability of one experimental condition *e* (e.g., the rat entering the left port) as

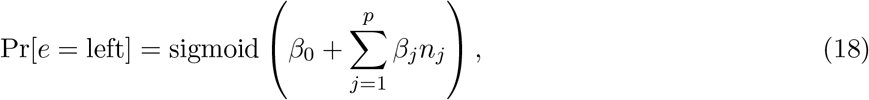

where *n*_*i*_ is the neural activity of the *i*th neuron. The corresponding objective function is the log-likelihood, or

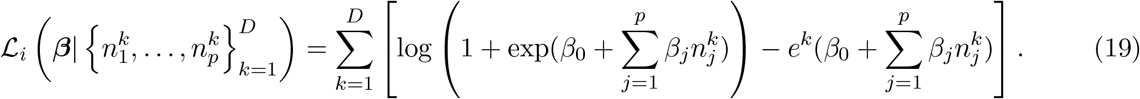

The baseline approach consisted of solving this objective function with an *ℓ*_1_ penalty. UoI_Logistic_ utilized objective function (19) with an *ℓ*_1_ penalty in the selection module, and Eq (19) alone in the estimation module.

#### Cross-validation, model training, and model testing

Each dataset was split into 10 folds after shuffling across samples (except for the basal ganglia data, which was split into 5 folds due to fewer samples). When appropriate, the folds were stratified to contain equal proportions of samples across experimental setting (e.g., stimulus value or behavioral condition). In each task, we fit 10 models (or five, for basal ganglia) by training each on 9 (4) folds, and using the last fold as a test set. Hyperparameter selection for baseline procedures was performed via cross-validation within the training set of 9 (4) folds. Meanwhile, all resampling for the UoI procedures was also performed within the training set. Model evaluation statistics (selection ratio, predictive performance, Bayesian information criterion) are reported as the median across the 10 models. Any measures that operate on the fitted models (e.g. coefficient value, network formation, modularity, etc.) were calculated by using the model that is formed by taking the median parameter value across folds.

#### Statistical tests

We used the Wilcoxon signed-rank test [79] to assess whether the distributions of selection ratios and predictive performances, across units, were significantly different between the UoI models and the baseline models. Importantly, we did not apply the test to the distribution of BICs, since differences in BIC are better interpreted as approximations to Bayes factors [80]. To assess whether distributions of UoI and baseline model parameters were significantly different, we used the Kolmogorov-Smirnov test. We applied a significance level of *α* = 0.01 for all statistical tests.

#### Effect size

To fully capture the difference in model evaluation metrics beyond statistical significance, we measure effect size using Cohen’s *d* [81]. For two groups of data with sample sizes *D*_1_, *D*_2_, means *μ*_1_, *μ*_2_, and standard deviations *s*_1_, *s*_2_, Cohen’s *d* is given by

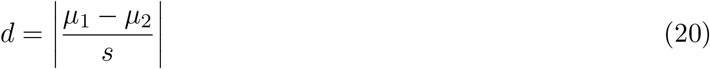

where *s* is the pooled standard deviation:

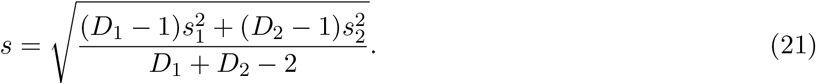

We often considered cases where *n*_1_ = *n*_2_, implying that 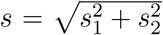. Values of *d* on the order of 0.01 indicate very small effect sizes, while *d* > 1 indicates a very large effect size [82].

### 2.6 Network creation and analysis

#### Network creation

We created directed graphs by filling the adjacency matrix *A*_*ij*_ with the coefficient *β*_*ij*_ (i.e., the coupling coefficient for neuron *j* in the coupling model for neuron *i*). Meanwhile, we created undirected networks from coupling models by symmetrizing coefficients [83]. Specifically, the symmetric adjacency matrix satisfies 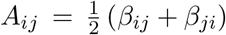 where *β*_*ij*_ is the coefficient specifying neuron *i*’s dependence on neuron *j*’s activity, and vice versa for *β*_*ji*_. Thus, the network lacked an edge between vertices (neurons) *i* and *j* if only if neuron *i*’s coupling model did not depend on neuron *j*, and neuron *j*’s coupling model did not depend on neuron *i*. This adjacency matrix is weighted in that each entry depends on the magnitudes of the coupling coefficients. However, we can also consider an unweighted, undirected graph, whose adjacency matrix is simply the binarization of *A*_*ij*_.

#### Modularity

The modularity *Q* is a scalar value that measures the degree to which a network is divided into communities [84]. We operate on the undirected graph described by the binarized adjacency matrix, described in the previous section. Suppose each vertex *v* is partitioned into one of *c* communities, where vertices within a community are more likely to be connected with each other than vertices between communities. Then, the modularity is defined as

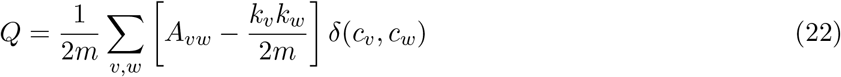

where *c*_*v*_ denotes community identity, *k*_*v*_ is the degree of vertex *v*, and *m* is the total number of edges. Thus, the modularity is greater than zero when there exist more edges between vertices within the same community than might be expected by chance according to the degree distribution. Specifically, *Q* is bounded within the range [−1/2, 1], where *Q* > 0 indicates the existence of community structure.

We calculated modularity with the Clauset-Newman-Moore greedy modularity maximization algorithm[85]. This procedure assigns vertices to communities by greedily maximizing the modularity, and then calculating *Q* using the ensuing community identities.

#### Small-worldness

Small-world networks are characterized by a high degree of clustering with a small characteristic path length [17, 86, 87]. There are multiple measures used to quantify the degree to which a network is small-world. We use *ω*, which can be expressed as

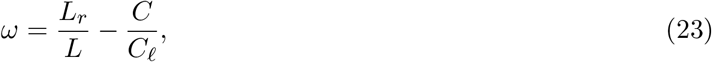

where *L* is the characteristic path length of the network, *L*_*r*_ is the characteristic path length for an equivalent random network, *C* is the clustering coefficient, and *C*_*ℓ*_ is the clustering coefficient of an equivalent lattice network [88]. The quantity *ω* is bounded within [−1, 1], where *ω* close to 0 indicates that the graph is small-world. When *ω* is close to 1, the graph is closer to a random graph, while *ω* close to −1 implies the graph is more similar to a lattice graph.

## 3 Results

Parametric models are ubiquitous data analysis tools in systems neuroscience. However, their usefulness in understanding a neural system hinges on the assumption that their parameters are accurately selected and estimated. By accurate selection, we mean low false positives and false negatives in setting parameters equal to zero; by accurate estimation, we mean low-bias and low-variance in the parameter estimates. The potential neuroscientific consequences of improper selection or estimation during inference are generally not well understood. Thus, we studied selection and estimation in common systems neuroscience models by comparing the properties of models inferred by standard methods to those inferred by the Union of Intersections (UoI) framework, which has been shown to achieve state-of-the-art inference in such models. We fit models spanning functional coupling (coupling networks from auditory cortex, V1, and M1), sensory encoding (spatio-temporal receptive fields from retinal recordings and tuning curves from auditory cortex), and behavioral decoding (classifying behavioral condition from basal ganglia recordings). We analyzed the fitted models to assess whether improvements in inference impact the resulting neuroscientific conclusions.

### 3.1 Highly sparse coupling models maintain predictive performance

Functional coupling models detail the statistical interactions between the constituent units (e.g., neurons, electrodes, etc.) of a population. Such models can be used to construct networks, whose structural properties may elucidate the functional and anatomical organization of the neurons within the population [14, 15, 19]. Enhanced sparsity in these models could result in different inferred functional sub-networks reflected in the ensuing graph. Furthermore, obtaining biased parameter estimates obscures the relative importance of neuronal relationships in specific sub-populations. Therefore, precise selection and estimation in coupling models is necessary to properly relate the network structure to the statistical relationships between neurons.

We examined the possibility of building highly sparse and predictive coupling networks by fitting coupling models to data from three brain regions: recordings from auditory cortex (AC), primary visual cortex (V1), and primary motor cortex (M1). The AC data consisted of micro-electrocorticography (*μ*ECoG) recordings from rat during the presentation of tone pips (Dougherty & Bouchard, 2019: DB). The V1 data consisted of single-unit recordings in macaque during the presentation of drifting gratings (Kohn & Smith, 2016: KS). The M1 data consisted of single-unit recordings in macaque during self-paced reaches on a grid of targets (O’Doherty, Cardoso, Makin, & Sabes: OCMS). See Methods for further details on experiments, model fitting, and metrics used for model evaluation, and see Table B.1 for a model statistic summary.

We constructed coupling models consisting of either a regularized linear model (AC) or Poisson model (V1, M1) in which the activity of a electrode/single-unit (i.e., node) was modeled using the activities of the remaining electrodes/single-units in the population. Thus, each dataset had as many models as there were distinct electrodes/single-units. We quantified the size of the fitted models with the selection ratio, or the fraction of parameters that were non-zero. We compare the selection ratio between baseline and UoI coupling models across electrodes/single-units in Fig 4a. For all three brain regions, UoI models exhibited a marked reduction in the number of utilized electrodes/single-units. Specifically, UoI models used 2.24 (AC), 2.21 (V1), and 5.50 (M1) times fewer features than the corresponding baseline models. Across the populations of electrodes/neurons, this reduction was statistically significant (*p* << 0.001; see Table B.1b) with large effect sizes (AC: *d* = 1.74; V1: *d* = 2.26; M1: *d* = 2.49; see Table B.1b). Interestingly, while the reduction in features for AC and V1 are roughly similar, the M1 models exhibit a much larger reduction in selection ratio, an observation that holds across the three M1 datasets. This disparity in feature reduction across brain regions indicates that false positives in the baseline coupling models may reflect differences in the nature of neural activity for these regions.

**Figure 4:**
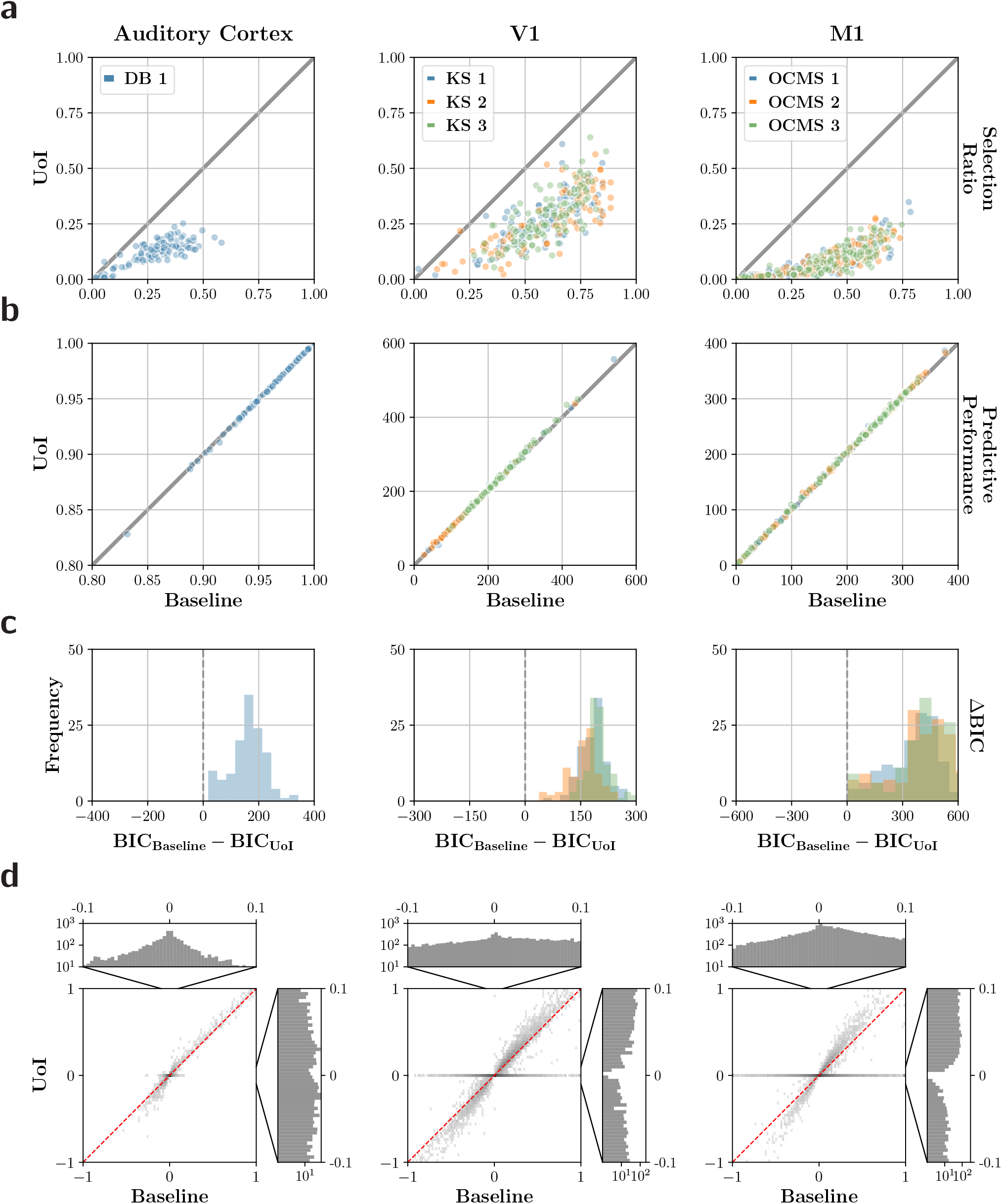
Highly sparse coupling models maintain predictive performance. **a-c.** Comparison of coupling models fit using baseline and UoI approaches to recordings from the auditory cortex (AC), primary visual cortex (V1), and primary motor cortex (M1). Each column corresponds to a different recording area (titles), and each row corresponds to a different model evaluation measure (right side labels). Colors denote different data sets within the same experiment, if available. **a.** The selection ratio, where the *y*-axis refers to the UoI model and *x*-axis the baseline model. Each point represents a coupling model for a specific single-unit/electrode. Gray line denotes the identity line. **b.** Comparison of predictive performance. **c.** The distribution of BIC differences across coupling models. **d.** Comparison of the coefficient values inferred by UoI (*y*-axis) and baseline (*x*-axis) methods. Data is depicted on a hexagonal 2D histogram with a log intensity scale. The marginal distributions of the non-zero coefficient values are shown on the top (baseline) and side (UoI) for each brain area. Note that the 1D histograms display the distribution in a more restricted domain than the 2D histograms, as depicted by the black lines.

We assessed whether the reduction in features resulted in meaningful loss of predictive accuracy. We measured predictive accuracy using the coefficient of determination (*R*^2^) for linear models (AC) and the deviance for Poisson models (M1, V1), both evaluated on held out data. Note that in contrast to *R*^2^, lower deviance is preferable. The predictive performances of baseline and UoI models for each brain region are compared in Fig 4b. We observed that there is almost no change in the predictive performance across brain regions, with most points lying on or close to the identity line. We note that while the differences in performance across all models were statistically significant (AC: *p* < 10^*−*3^; V1: *p* << 0.001; M1: *p* << 0.001; see Table B.1c), the effect sizes of the reduction in predictive performance were very small (AC: *d* = 0.005; V1: *d* = 0.05; M1: *d* = 0.03; see Table B.1c), making it irrelevant in practice. Thus, these results imply that highly sparse coupling methods exhibit little to no loss in predictive performance across brain regions and datasets.

We captured the two previous observations — increased sparsity and maintenance of predictive accuracy — with difference in Bayesian information criterion (BIC) between baseline and UoI methods, ΔBIC = BIC_baseline_ − BIC_UoI_. Lower BIC is preferable, so that positive ΔBIC indicates that UoI is the more parsimonious and preferred model. The distribution of ΔBIC across coupling models is depicted in Fig 4c. ΔBIC is positive for all models, with a large median difference (AC: 170; V1: 149; M1: 186; see Table B.1d), providing very strong evidence against the baseline models.

To characterize the functional relationships inferred by the coupling models, we examined the distribution of coefficient values. We normalized each model’s coefficients by the coefficient with largest magnitude across the baseline and UoI models, and concatenated coefficients across models and datasets. We visualized the baseline and UoI coefficient values using a 2-d hexagonal histogram (Fig 4d). First, we observed a density of bins above (positive coefficients) and below (negative coefficients) the identity line (Fig 4d: red dashed line). This indicates that the magnitude of non-zero coefficients as fit by UoI are larger than the corresponding non-zero coefficient as fit by the baseline, demonstrating the amelioration of shrinkage and therefore reduction in bias. Next, we observed a density of bins on the *x* = 0 line, indicating a sizeable fraction of coefficients determined to be non-zero by baseline methods are set equal to zero by UoI. This density corroborates the reduction in selection ratio observed in Fig 4a. We further note that the density of bins on the *x* = 0 line encompass a wide range of baseline coefficients values, especially for the V1 and M1 datasets. This implies that utilizing a thresholding scheme (e.g., BoATS) based on the magnitude of the fitted parameters for a feature selection procedure will not reproduce these results. Lastly, we observe no density of bins along the *y* = 0 line, which indicates that UoI models are likely not identifying the existence of functional relationships which do not exist (i.e., suffering from false positives).

While many of the coefficients set equal to zero by UoI have large magnitude (as measured by baseline methods), the bulk of density lies in coefficients with small magnitude. We found that the difference in distributions of non-zero coefficients between the two procedures is statistically significant (*p* << 0.001; Kolmogorov-Smirnov test). We highlight the marginal distribution of non-zero coefficients whose magnitudes are small (Fig 4d, top and side histograms). While the baseline histograms (Fig 4d, top histograms) have the largest density of coefficients close to zero, the UoI histograms, in a similar range, exhibit a large reduction in density. Together, these results indicate that standard inference methods produce overly dense networks with highly biased parameter estimates, giving rise to qualitatively different parameter distributions.

#### Accurate inference enhance visualization, increase modularity, and decrease small-worldness in functional coupling networks

Functional coupling networks are useful in that they provide opportunities to visualize the statistical relationships within a population. Furthermore, their graph structures can be analyzed to characterize global properties of the network. The previous results show that improved inference gives rise to equally predictive models, but with much greater sparsity and qualitatively different parameter distributions. Thus, we next determined the impact on network visualization and structure. To this end, we constructed and analyzed both directed networks from the coupling coefficients and undirected networks by symmetrizing coupling coefficient values (see Methods).

We first visualized the AC networks by plotting the baseline and UoI networks according to their spatial organization on the *μ*ECoG grid (Fig 5a). Each vertex in Fig 5a is color-coded by preferred frequency, while the symmetrized coupling coefficients are indicated by the color (sign) and weight (magnitude) of edges between vertices. We observed that the UoI network is easier to visualize, with densities of edges clearly demarcating regions of auditory cortex. This is contrast to the baseline network, whose lack of sparsity makes it difficult to extract any meaningful structure from the visualization. For example, the UoI network exhibits a clear increase in edge density in primary auditory cortex (PAC) relative to the posterior auditory field (PAF) and ventral auditory field (VAF). Thus, the increased sparsity in UoI networks reveals graph structure that ties in closely with general anatomical structure of the recorded region.

**Figure 5:**
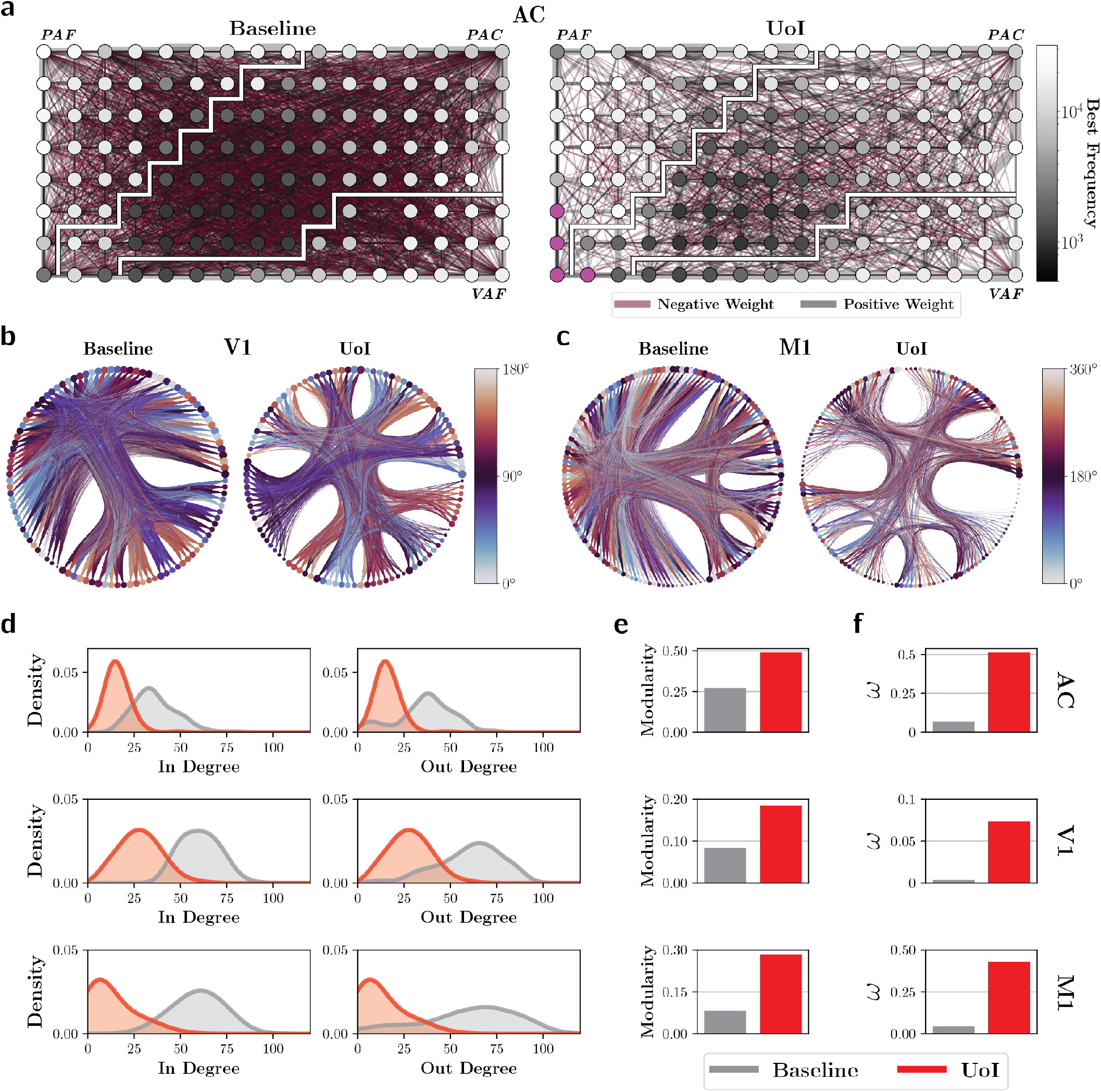
Accurate inference enhances visualization, increase modularity, and decrease small-worldness in functional coupling networks. **a.** Networks obtained from auditory cortex data. Vertices are organized according to their position on the electrocorticography grid. Vertices are color coded by preferred frequency, with fuchsia vertices denoting non-tuned electrodes. Edge width increases monotonically with edge weight while edge color denotes the sign of the weight. White lines segment the grid according to the regions of auditory cortex. **b.** Visualization of example coupling networks for visual cortex recordings, with vertices color-coded by preferred tuning. Edges are bundled according to detected communities, while vertex size corresponds to its degree. Note that stimulus encoding is cyclical (static grating angle from 0° to 180°). **c.** Visualization of coupling networks for motor cortex recordings, with vertices color-coded by preferred tuning. Edges are bundled according to detected communities, and vertex size corresponds to its degree. Note that stimulus encoding is cyclical (movement direction angle from 0° to 360°). **d-f.** Comparison of common graph metrics evaluated on UoI networks (red) and baseline networks (gray). Each row corresponds to a distinct brain region (left: AC, V1, and M1 from top to bottom). **d.** In-degree and out-degree densities. **e.** The graph modularity, averaged across datasets. **f.** The small-worldness as quantified by *ω*, and averaged across datasets.

We visualized the V1 and M1 networks by fitting nested stochastic block models to the directed graphs and plotting the ensuing structure in a circular layout with edge bundling (Fig 5b, c). The nested stochastic block model identifies communities of vertices (neurons) in a hierarchical manner. We color-coded the vertices of the visualized graphs according to preferred tuning (drifting grating angle for V1 networks and hand movement angle for M1 networks). The UoI V1 network exhibits clear structure, with specific communities identifying similarly tuned neurons (Fig 5b). The baseline V1 network does not exhibit as clear structure, and was highly unbalanced, with more than half the neurons placed in the same community. For the M1 data, the UoI communities were more balanced, though they lacked clear association with tuning properties. Together, these results demonstrate that enhanced sparsity of functional coupling networks facilitate their interpretation through cleaner visualizations and a clearer connection to functional response properties.

The plots above suggest different graph structures in the networks extracted by UoI and baseline methods. Thus, we first calculated the in-degree and out-degree distributions of the vertices in both networks (Fig 5d). We observed that the in-degree and out-degree distributions for the UoI networks are much smaller, as one might expect due to the reduction in edges. Furthermore, the in- and out-degree distributions describing the UoI networks are similar, in contrast to those of the baseline networks. Next, we calculated the modularity of the networks, which quantifies the degree to which the networks exhibit community-like structure. We found that the modularity for the UoI networks is much larger than that of baseline networks, indicating that UoI networks express more community structure than baseline networks (Fig 5e). These results corroborate the visual findings in Fig 5b. Since modularity implicitly depends on degree distribution, the enhanced community structure exhibited by the UoI networks is not simply a property of the reduction of in- and out-degrees, emphasizing that the enhanced sparsity is functionally meaningful. We found similar findings in functional networks built from linear models, rather than Poisson models, implying that more accurate inference ensures that the structure of coupling networks persists across the type of underlying model (Appendix A and Fig. A).

Finally, we examined the small-worldness of the networks, a graph structure commonly used to describe brain networks. Small-world networks are characterized by a high degree of clustering and low characteristic path length, making them efficient for communication. We used *ω* to calculate small-worldness, whose values are bounded by the range [−1, 1], with *ω* close to −1, 0, and 1 indicative of lattice, small-world, and random structure, respectively. Interestingly, UoI networks are considerably less small-world than the baseline networks (Fig 5f). However, we note that all networks are more small-world than they are random. Furthermore, the small-worldness of the networks is dependent on the brain region. For example, the V1 networks exhibit substantially more small-worldness than the auditory cortex or M1 networks. Together, these results demonstrate that several properties of networks can be substantially altered by the utilized inference procedure, and that UoI networks are more modular and less small-world.

### 3.2 Parsimonious tuning from encoding models

A long-standing goal of neuroscience is to understand how the activity of individual neurons are modulated by factors in the external world (e.g., how the position of a moving bar is encoded by a neuron in the retina). In such encoding models, incorrect feature selection or parameter bias may mistakenly implicate factors in the production of neural activity, or misstate their relative importance. Thus, we examined how improved inference impacts tuning models, where an external stimulus is mapped to the corresponding evoked neural activity.

We first fit spatio-temporal receptive fields (STRFs) to single-unit recordings from isolated mouse retinal ganglion cells during the presentation of a flicking black or white bar stimulus (generated by a pseudo-random sequence). We used a linear model with a lasso penalty to fit STRFs (i.e., regularized, whitened spike-triggered averaging) to recordings from 23 different cells, using a time window of 400 ms. Thus, the fitted STRFs were two dimensional, with one dimensional capturing space (location in the bar stimulus) and the other capturing the time relative to neural spiking. For further experimental and model fitting details, see Methods. See Table B.2 for a dataset and model statistic summary.

The fitted STRFs for an example retinal ganglion cell are depicted in Fig 6a. The UoI STRF captures the ON-OFF structure exhibited by the baseline STRF. However, the UoI model is noticeably sparser, resulting in a tighter spatial receptive field. The features set to zero by UoI (relative to baseline) include regions both further from the dominant ON-OFF structure, and regions very close to the center. Additionally, the coefficient values of the UoI STRF are noticeably larger in magnitude, implying that the improved estimation has alleviated shrinkage. These results suggest that the baseline procedure may produce false positives in both the central and distal regions of the STRF, implying that the relevant regions for predicting the activity of retinal ganglion cells may be more restricted.

**Figure 6:**
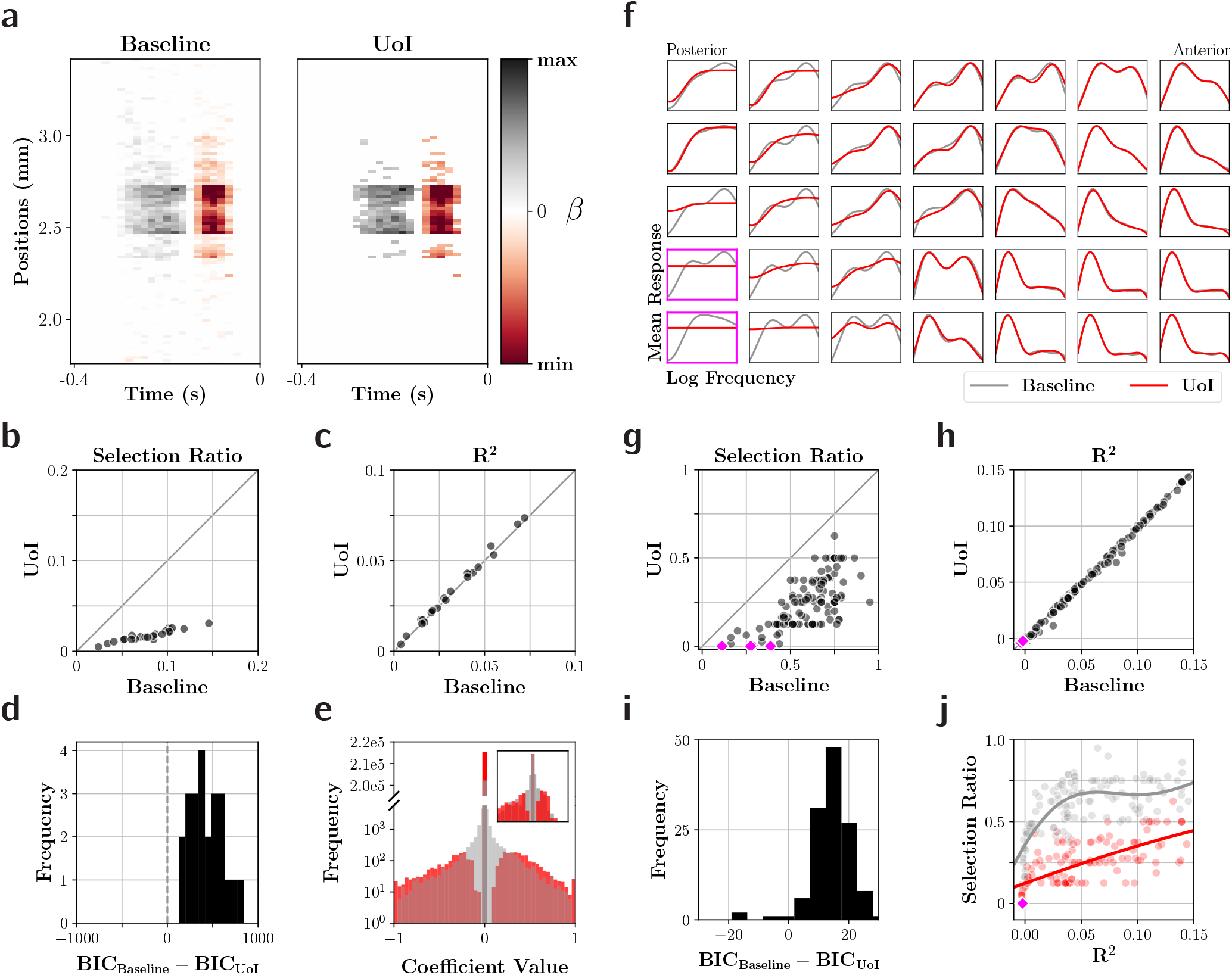
Parsimonious tuning from encoding models. **a-e.** Analysis of spatiotemporal receptive fields (STRFs) fit to spikes from mouse retinal ganglion cells using a 1d white noise stimulus. **a.** Example STRF extracted by baseline (left) and UoI (right) procedures. **b-c.** Quantitative comparison of STRFs extracted with UoI (*y*-axis) and baseline (*x*-axis) procedures. Each point represents a unique STRF. Gray line denotes identity. **b.** Comparison of selection ratios for each STRF. **c.** Comparison of coefficient of determination (*R*^2^) on held out data. **d.** The distribution in BIC differences across STRFs. Dashed line denotes equal BIC, i.e. ΔBIC = 0. **e.** Distribution of normalized coefficient values across all baseline (gray) and UoI (red) STRFs. The broken *y*-axis is log-scale below the break and a regular scale above the break. Inset shows distribution of coefficients for STRFs depicted in **a. f-j.** Analysis of tuning curves extracted from micro-electrocorticography recordings on rat auditory cortex during the presentation of tone pips. **f.** Examples of tuning curves for a subset of electrodes on the grid, as fit by baseline (gray) and UoI (red) procedures. Pink outlines denote tuning curves set exactly equal to zero by UoI. **g-h.** Quantitative comparison of tuning curves extracted with UoI (*y*-axis) and baseline (*x*-axis) procedures. Panels are structured similarly as panels **b-c.**. Pink points highlight tuning curves that had a selection ratio of zero, as determined by UoI. **i.** The distribution in BIC differences across electrode tuning curves, structured similarly as panel **d. j.** Selection ratio plotted against coefficient of determination for baseline (gray) and UoI (red) procedures. Each point denotes a STRF. Trendlines are fit with Gaussian process regression.

We compared the selection ratios across fitted STRFs (Fig 6b) and found UoI fits to be substantially sparser, with a median reduction factor of 4.98. This reduction was statistically significant (*p* << 0.001; see Table B.2b) and had a very large effect size (*d* = 3.05; see Table B.2b). At the same time, UoI models exhibited a statistically significant improvement in *R*^2^ (*p* < 0.01; see Table B.2c), but with a very small effect size, making the improvement irrelevant in practice (*d* = 0.05; see Table B.2c). Meanwhile, the BIC differences (Fig 6d) were all large and positive (median difference = 654; see Table B.2d), providing very strong evidence in favor of the UoI model. Lastly, we compared the distribution of baseline and UoI encoding coefficients, normalized to the largest magnitude coefficient. We found evidence that the UoI models exhibited reduced shrinkage (Fig 6e: larger tails), and a substantial reduction in false positives (Fig 6e: reduced density at origin). Thus, improved inference resulted in STRFs with tighter structure, in better agreement with theoretical work characterizing such receptive fields and more accurately reflecting the visual features that explain the production of neural activity in retinal ganglion cells [89, 90].

Across neurons, we observed selection ratios and predictive performances spanning a wide range of values (Fig 6b, c). We might expect that models with little predictive accuracy utilize fewer features, since poor predictive performance indicates that the provided features are inadequate. For example, in the limit that the model has no predictive capacity (*R*^2^ ≤ 0), the model should utilize no features, since such an *R*^2^ indicates that none of the available features are relevant for reproducing the response statistics better than the mean response value. Therefore, we sought to determine whether inaccurate inference mistakenly identifies models as tuned (i.e., non-zero tuning features), when in fact a “non-tuned” model is appropriate (e.g., an intercept model: all features set equal to zero). To this end, we utilized a dataset in which the feature space dimensionality is small. This scenario provides a suitable test bed for assessing whether an intercept model could arise, and if such a model is appropriate given the response statistics of the data. We examined a dataset consisting of *μ*ECoG recordings from rat auditory cortex during the presentation of tone pips at various frequencies. We employed a linear tuning model mapping frequency to the peak (*z*-scored relative to baseline, see Methods), high-*γ* band analytic amplitude of each electrode. The model features consisted of 8 Gaussian basis functions that tiled the log-frequency space.

We first examined whether more accurate inference resulted in any qualitative changes in the fitted encoding models. We plotted the fitted tuning curves as a function of log-frequency for a subset of electrodes arranged according to their location on the *μ*ECoG grid (Fig 6f). Interestingly, the baseline and UoI tuning curves exhibit similar structure for a large fraction of the electrodes on the grid, in many cases matching closely (e.g., Fig 6f: anterior side of grid). In other cases, particularly on the posterior side of the grid, the UoI tuning curves exhibit similar broad structure with noticeable smoothing, indicating that the improved inference has simplified the tuning model.

We compared the selection ratio of the models (Fig 6g), finding that the UoI tuning curves utilize fewer features than those fit by baseline, with a median reduction factor of 2.5 that is statistically significant (*p* 0.001; see Table B.2b) and a large effect size (*d* = 2.19; see Table B.2b). Furthermore, despite a statistically significant decrease in *R*^2^ (Fig 6h) across electrodes (*p* 0.001; see Table B.2c), the observed effect size is very small (median Δ*R*^2^ = 0.001; *d* = 0.05; see Table B.2c). Meanwhile, we observed a median BIC difference of 19.4, providing very strong evidence in favor of the UoI models (Fig 6i; see Table B.2d). Taken together, these results imply that the reduction in features did not harm the predictive performance of the tuning models, thereby enhancing their parsimony. We highlight four electrodes whose selection ratios, according to UoI, are exactly zero in Fig 6g (pink points). These four “non-tuned” electrodes are among the least predictive, with *R*^2^ close to or below zero for both baseline and UoI methods (Fig 6h: pink points). Interestingly, the baseline selection ratio for one of these models was close to 0.4, indicating that UoI is not trivially generating intercept models. We visually examined the frequency-response areas (FRAs) of these four electrodes, which detail the mean response values as a function of sound frequency and amplitude (Fig. C). We compared them to the FRAs of two randomly chosen electrodes, finding they had little discernible structure relative to the “tuned” FRAs. Thus, while more accurate inference in encoding models may not always result in perceptible changes in their appearance (e.g., Fig 6f), there are cases where inaccurate inference may mistakenly imply that a constituent unit is tuned, when in fact the stimulus features are not relevant for capturing its response statistics. To understand the behavior of the selection ratio as *R*^2^ → 0, we examined the relationship between selection ratio and *R*^2^ for baseline (gray) and UoI (red) models (Fig 6j). We found that the sparser models exhibit lower predictive power, with model trends predicting that at *R*^2^ = 0, the selection ratio for the baseline model will be 0.35 ± 0.10 (mean ± 1 s.d.) while the UoI selection ratio will be 0.12 ± 0.10. This demonstrates that baseline procedures can suffer from false positives even when their fitted models exhibit little to no explanatory power. Overall, our results reveal that improved inference more accurately identifies units as non-tuned when their encoding models lack predictive ability.

### 3.3 Decoding behavioral condition from neural activity with a small number of single-units

Decoding models describe which neuronal sub-populations contain information relevant for an external factor, such as a stimulus or a behavioral feature. Such models can identify which neurons may be useful to a downstream population for a task that requires knowledge of an external factor. Specifically, a decoding model’s non-zero parameters can be interpreted as the sub-population of neurons containing the task-relevant information, emphasizing the need for precise selection. Additionally, the model details how specific neurons describe the decoded variable through the magnitudes of its parameters, requiring unbiased estimation. Thus, we sought to assess the degree to which accurate inference impacts data-driven discovery in neural decoding models.

For this analysis we examined 54 single units in the rat basal ganglia (18 units from globus pallidus pars externa, GPe, and 36 from the substantia nigra pars reticulata, SNr) that were recorded simultaneously during performance of a behavioral task involving rapid leftward or rightward head movements in response to cues. Details of the task are given in [72]; the analysis was restricted to trials in which a correct head movement was made. Thus, the decoding model consisted of binary logistic regression predicting trial outcome (left or right) using the single-unit spike counts as features. We fit the logistic regression with an *ℓ*_1_ penalty (baseline) and the UoI framework (UoI_Logistic_). Further details on the experimental setup and model fitting can be found in Methods. See Table B.3 for a summary of the dataset and fitted model statistics.

The selection ratios for GPe and SNr, as obtained by baseline and UoI procedures, are depicted in Fig 7a. In GPe, the UoI decoding models utilized about half the number of parameters as the baseline procedures. Meanwhile, in SNr, the UoI models utilized four times fewer parameters than the baseline. Furthermore, we observe that UoI model sizes were more consistent across folds of the data. For example, the SNr decoding models estimated anywhere from 1 to 21 parameters (out of 36) depending on the data fold, while UoI models consistently used only 2 or 3 parameters (Fig 7a: IQR bars). These results validate that neural decoding models are capable of utilizing fewer features to predict relevant behavioral features. Furthermore, the stability principle ensures that these features are more robust to perturbations of the data (e.g., random subsamples).

**Figure 7:**
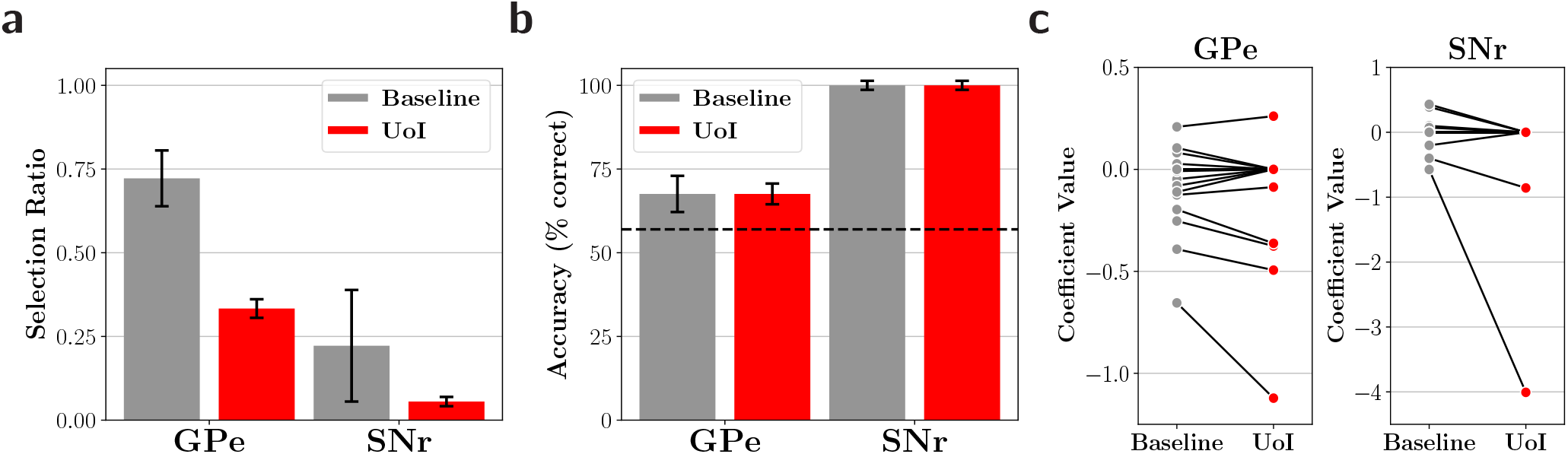
Behavioral condition can be decoded with a small number of single-units at no loss in accuracy. Decoding models were applied to single-unit recordings from rat basal ganglia. The models, consisting of binary logistic regression, predicted whether the rat went left or right on a stop signal task. Single-unit recordings were used from the globus pallidus pars externa (GPe: 18 units) and the substantia nigra pars reticulata (SNr: 36 units). **a-b.** Evaluation of decoding models fit via the UoI (red) and baseline (gray) procedures. Bar heights indicate median across five data folds, while error bars denote IQR. **a.** Comparison of the selection ratios. **b.** Comparison of left/right classification accuracy on held-out data. Dashed line denotes accuracy by chance. **c.** Comparison of fitted coefficient values extracted by baseline and UoI procedures in GPe and SNr. The decoding models fit by UoI utilize about 3 times fewer single-units at no cost to accuracy, indicating that task relevant information is contained in a small number of single-units.

To examine whether the use of fewer single-units decreased predictive performance, we evaluated the decoding models’ classification accuracy on held-out data, depicted in Fig 7b. The classification accuracy of the UoI models is equal to that of the baseline models for both GPe (67%) and SNr (100%). Furthermore, in both regions, the classification accuracy is greater than chance (56%), implying that the models are extracting meaningful information about the behavioral condition from the neural response. Interestingly, the median SNr models achieve perfect classification accuracy. The UoI model achieves this performance utilizing only 2 neurons, in contrast to median baseline model, which utilizes 8. Therefore, the activities of only a small subset of neurons are required to predict the behavioral condition, an observation that required accurate inference to consistently capture.

We examined the fitted coefficient values for each brain region and fitting procedure (Fig 7c). First, we observed that all coefficients set equal to zero by the baseline procedure are also set equal to zero by UoI. Additionally, the coefficients set equal to zero by UoI, but not the baseline procedure, typically have smaller magnitudes than the coefficients that are non-zero for both procedures. Finally, the coefficients set non-zero by UoI have larger magnitudes relative to their value under the baseline procedure. These observations imply that the UoI procedure, for this task, consistently utilized only the most important neurons to predict the behavioral condition. At the same time, UoI elevated the coefficient values relative to the baseline procedure, implying that baseline procedures suffered from substantial parameter shrinkage. Overall, these results demonstrate that task-relevant information is conveyed by a sparse subset of basal ganglia neurons, especially in SNr. SNr is a basal ganglia output nucleus, receiving converging inputs from multiple basal ganglia structures including GPe. The finding that SNr decodes behavioral output more selectively and accurately compared to GPe is consistent with the idea that SNr is closer to post-decision behavioral outputs, whereas GPe represent internal preparatory states [72].

## 4 Discussion

Parametric models are used pervasively in systems neuroscience to characterize neural activity. The parameters of these models must be precisely selected and estimated in order to ensure accurate interpretation, particularly in the sparse parameter regime that is desirable for neural data. Here, we used the UoI framework, which achieves state-of-the-art performance in balancing selection and estimation, to assess the degree to which poor parameter inference may impact neuroscientific interpretation of parametric models. We fit functional coupling, encoding, and decoding models to a battery of neural datasets, using standard and UoI inference procedures. We found that, across all models, the number of non-zero parameters could be reduced by a factor of 2–5, while maintaining predictive performance. Furthermore, we found broader, structural differences in the models beyond enhanced sparsity, which resulted in concrete changes to their neuroscientific interpretation.

The parameters obtained in coupling models denote the existence and strength of functional relationships between constituent units in a population. We observed striking differences in the distribution of these parameter estimates, which impacted the graph structure, manifesting in increased modularity and decreased small-worldness. These results do not directly contradict previous work characterizing brain networks as small-world, but do reduce the magnitude of the characterization [17, 87, 91] (though see [92]). Coupling model parameters have also been assessed for their recapitulation of synaptic weight distributions in neural circuits, in some cases identifying parameter biases induced by specific dynamical regimes of neural activity [93]. The coupling parameters extracted by UoI better reflect the weight distribution as observed in neural circuits, which is characterized by sparse connectivity with a heavy-tailed distribution [94, 95]. This was not achieved by baseline procedures, suggesting the previously identified biases could instead be due to inaccurate inference. The salient differences in the inferred coupling parameter distributions we observed motivates similar examination in models that capture neural dynamics, such as vector auto-regressive models [96, 97], which could be further assessed by controllability metrics used in recent work on fMRI networks [98].

The parameters in encoding models detail which features modulate neural activity. We observed that the application of UoI to the encoding models highlighted cases where the fitted model had only zero parameters, other than the intercept. Such an intercept model implies that a tuning model may be inappropriate for capturing the response statistics of the constituent units. This observation can be understood as a natural consequence of stability enforcement during parameter inference: UoI benefits from the stability principle by only utilizing selected features that persist across data resamples. The use of data resamples mimics perturbing the dataset, ensuring that features are included only if they are robust to those perturbations. Thus, the stability principle enforces model parsimony by encouraging the use of fewer features, eliminating those that offer no predictive accuracy throughout the resamples. However, UoI prioritizes predictive accuracy in the model averaging step. Therefore, models will only be made “as simple as possible” (e.g., removing all features) when they possesses no predictive ability. In contrast to the auditory cortex, we fit spatio-temporal receptive fields on the retinal ganglion cells with typical ON-OFF structure [8]. However, the stability enforcement of UoI resulted in models that were more spatially constrained, in better agreement with theoretical work characterizing the receptive fields [90, 99]. More broadly, these results indicate that such improvements in parameter inference could serve to close the gap between experiment and theory in systems neuroscience.

Decoding models can inform which internal factors contain information about a task-relevant external factor. We found that decoding models could be fit using fewer single-units, at no cost to classification accuracy, implying that task-relevant information can be confined to a very small fraction of a neural circuit. This has implications for communication between brain areas: wiring constraints often restrict information transmission through a relatively smaller number of projection neurons, suggesting that these neurons contain the relevant information required to “decode” a given signal [100, 101]. These results, in which we identified a very small fraction of the neurons capable of accurate decoding, raise the possibility that these selected neurons are, in fact, the projection neurons. Decoding using fewer single-units also has practical implications. Furthermore, an abundance of work has considered the impact of correlated trial-to-trial variability on the fidelity of a neural code by assessing the decoding ability of neural populations as a function of population size [102]. These decoding analyses can be informed by knowledge of the sparse sub-populations predictive of an external factor, which these results indicate are smaller than previously thought. Brain-machine interfaces (BMIs), which rely on accurate decoding from neural activity to operate, could reduce their power consumption by using a decoder relying on fewer single-units. Together, these results imply that accurate inference procedures, more capable of discovering specific task-relevant neuronal sub-populations, could drive the development of normative theories of neural communication and decoding.

Across brain regions and models, UoI resulted in more parsimonious models with differences in predictive performance that were irrelevant in practice, as measured by Cohen’s *d*. However, statistical tests comparing the distribution of predictive performance between the baseline and UoI models revealed a statistically significant decrease in predictive performance for some cases (coupling models, AC tuning) and statistically significant increase in others (retinal STRF, decoding). Depending on one’s goals, relying solely on predictive performance to judge a model may be unreliable [75–77, 103]. In particular, because model interpretability depends crucially on the included features and their estimates, we prioritized feature selection and estimation. Cross-validated predictive accuracy is often a poor criterion for accurate feature selection. In these cases, the BIC, which captures model parsimony, serves as a more suitable criterion [46], and universally favored the UoI models (though we note there is no single preferred model selection criterion [46, 77, 104–106]).

We considered models of neural activity exclusively in terms of coupling, encoding, or decoding. However, past studies have built parametric models of neural activity by using other features or model structures. For example, the combination of coupling and encoding in a single model is a natural extension which has been examined extensively in previous work [6–8, 13, 107]. Other features that are not constrained within coupling, encoding, or decoding — such as spike-time history or global fluctuations — are also important to incorporate [7, 9, 108]. Additionally, latent variable models have been used to great success to capture, in particular, the dynamics of neural activity [109–111]. In this work, the stability principles used by UoI resulted in a significant difference in the model sparsity and estimated parameter distribution, which impacted interpretation. It is worthwhile to assess whether similar results can be achieved in these extended models, and if so, determine the neuroscientific consequences.

We restricted our analysis to generalized linear models, because their structure lends itself well to interpretation, making them ubiquitous in neuroscience and biology. However, the improvements we obtained by encouraging stability and sparsity in these models may extend to other classes of models. For example, dimensionality reduction methods have also played an important role in systems neuroscience [112]. The UoI framework is naturally extendable to such methods, including column subset selection [53] and non-negative matrix factorization [54]. Furthermore, recent work has found success in using artificial neural networks (ANNs) to model neural activity, which excel at predictive performance [113]. Since ANNs are highly parameterized, these models are not interpretable in the sense that their parameter values do not directly convey neuroscientific meaning. Instead, these models are often interpreted through the lens of learned representations or recapitulation of emergent properties of neural activity. Future work could assess whether the modeling principles explored in this work could have similar effects for ANNs modeling neural activity, especially given recent advancements in compressing such models [114].

In this work, using the UoI framework, we assessed the consequences of precise parameter inference specifically on systems neuroscience models. We used UoI because of its demonstrated success at parameter selection and estimation and our familiarity with the framework, but it is not alone in the class of inference algorithms that excel at parameter inference [60, 115]. Furthermore, model-based approaches to data-driven discovery are employed prolifically throughout biology [116], spanning genomics [117], cell biology [118], epidemiology [119], ecology [120] and others. Thus, the sparsity and stability properties invoked by the UoI framework, and other approaches, could serve to reshape model interpretability across a wide range of biological contexts.

## Acknowledgments

We thank the members of the Neural Systems and Data Science lab for helpful feedback and discussion. P.S.S. was supported by the Department of Defense (DoD) through the National Defense Science & Engineering Graduate Fellowship (NDSEG) Program. J.A.L. and K.E.B. were supported through the Lawrence Berkeley National Laboratory-internal LDRD “Deep Learning for Science” led by Prabhat. B.G. and J.D.B. were supported by NIH grants R01MH101697, R01NS078435 and R01DA045783, and the University of California, San Francisco.

## A Comparison of Poisson and linear coupling models for single-unit activity

We used a Poisson distribution to model single-unit spike count activity in the M1 and V1 datasets. However, past work has modeled single-unit activity with a linear-Gaussian model, after applying variance-stabilizing square root transform to the spike count responses. The degree to which a linear model can capture the functional relationships identified by a Poisson model for spiking data is unclear. Thus, we sought to characterize this capability, and its dependence on the inference procedure. We modeled the neural activity after a square-root transform 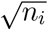 using a linear model

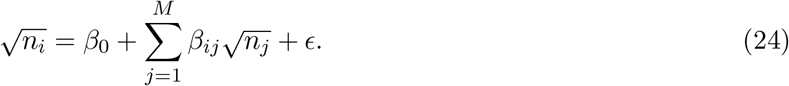

We fit this model with lasso optimization by coordinate descent (baseline) and UoI_Lasso_.

We compared the fitted selection profiles, i.e. the set *S* = {*i*|*β*_*i*_ ≠ 0} between the linear and Poisson models. To do so, we used the hypergeometric distribution, which describes the probability that *k* objects with a particular feature are drawn from a population of size *M* that has *K* total objects with that feature, using *m* draws without replacement. To frame this in terms of selection, suppose the Poisson model has |*S*_Poisson_| = *K* non-zero parameters out of the *M* possible features. Then, the probability the linear model, which has |*S*_linear_| = *m* non-zero parameters, would match the Poisson model on *k* such features is given by the hypergeometric distribution:

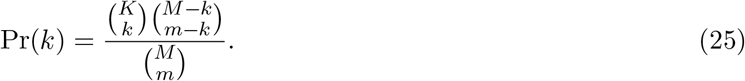

**Figure A:**
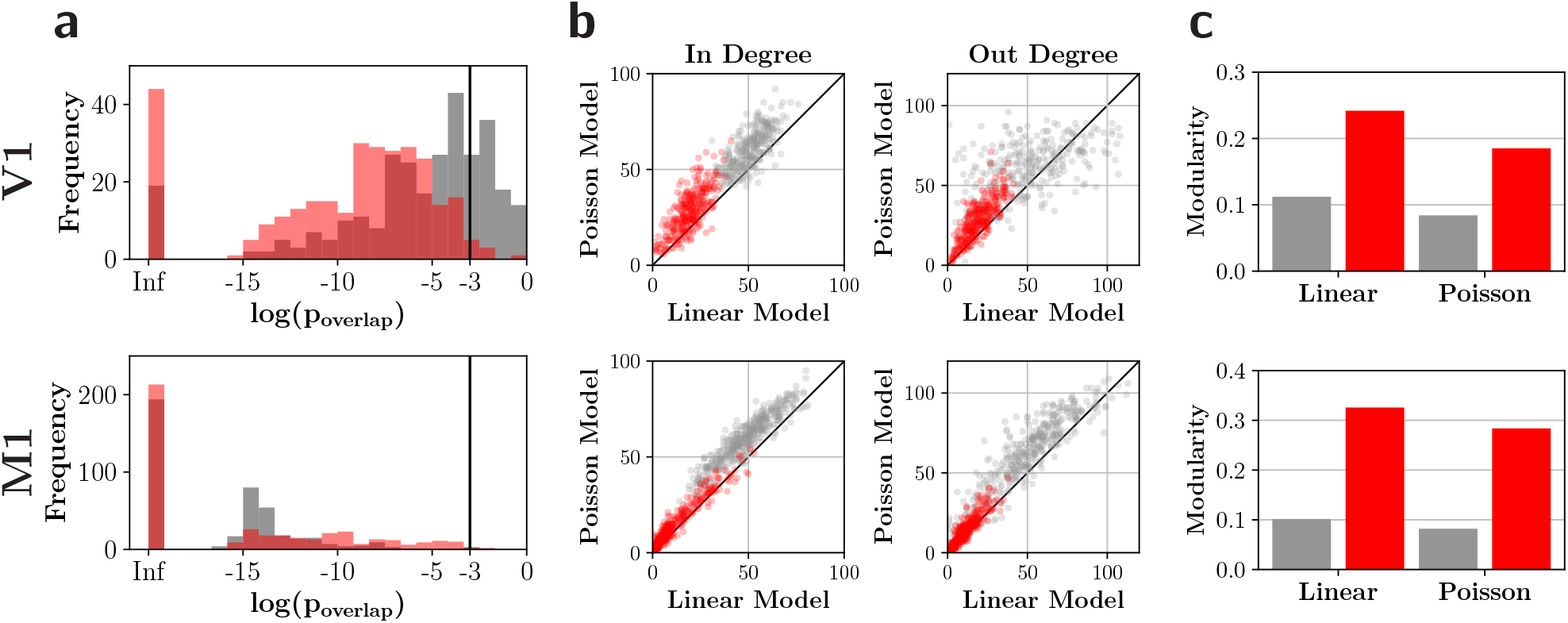
More accurate inference ensures that the structure of fitted coupling networks persists across the type of underlying model. Linear and Poisson coupling models were fit to the datasets with single-unit recordings (V1 and M1) using baseline (gray) and UoI (red) procedures. Top row corresponds to results from networks fit to V1 recordings, while bottom row corresponds to networks fit to M1 recordings. **a.** The (log) probability distribution of extracting a support (set of non-zero parameters) by the linear model that matches with the Poisson model support, according to a hypergeometric distribution. Vertical line denotes a p-value of 0.001 **b.** Comparison of the in-degree and out-degree distributions between the Poisson network (*y*-axis) and linear network (*x*-axis). Each point represents a single unit. Black line denotes identity. **c.** Graph modularity of linear and Poisson networks.

Thus, the probability that the selection profile would overlap at most as well by chance as the linear model is given by *p*_overlap_ = 1 − Pr(*k* < *k*_linear_). We compared the distribution of *p*_overlap_ across coupling models, calculated for both baseline and UoI procedures (Fig. A, panel a). For the V1 data, the UoI linear models better reproduced the Poisson selection profiles, with 98% of the profiles fit by UoI satisfying *p*_overlap_ < 0.001, compared to only 74% of the baseline selection profiles. In contrast, in the M1 data, both inference procedures fit linear models that closely matched the selection profiles of the Poisson models, with 99% of the selection profiles satisfying *p*_overlap_ < 0.001. Therefore, improved inference results in more consistent selection across models, and furthermore this consistency depends on brain region.

We constructed networks from the linear models as described in Methods, and calculated their in-degree and out-degree distributions. We compared the in-degree and out-degree distributions of the linear networks to the Poisson networks, finding a closer correspondence between the UoI models than for the baseline models in most cases (Fig. A, panel b). Specifically, the in-degrees of UoI models had a correlation of 0.742 (V1) and 0.969 (M1) compared to 0.717 (V1) and 0.924 (M1) for the baseline procedures. Similarly, we obtained out-degree correlations of 0.806 (UoI) and 0.531 (baseline) for V1 and 0.918 (UoI) and 0.924 (baseline) for M1. Lastly, we found that the UoI linear networks were more modular than the baseline linear networks. Interestingly, both were more modular than their Poisson counterparts (C, panel c). Taken together, these results imply that a more precise inference framework better preserves structure across model types.

## B Supplementary Tables

### B.1 Functional coupling dataset summary and fitted model statistics

Dataset summary (Table B.1a) provides details on the datasets used to fit functional coupling models, including the number of units and samples across brain region and recording session. The following tables provide statistics summarizing aspects of the fitted baseline and UoI models, including selection ratio (Table B.1b), predictive performance (Table B.1c), and Bayesian information criterion (Table B.1d).

**Table B.1a:** Dataset summary

| Brain Region | Number of Units | Number of Samples |
| --- | --- | --- |
| AC | 125 | 4200 |
| VC 1 | 106 | 2400 |
| VC 2 | 88 | 2400 |
| VC 3 | 112 | 2400 |
| MC 1 | 136 | 4089 |
| MC 2 | 146 | 4767 |
| MC 3 | 147 | 4400 |

**Table B.1b:** Selection ratio

| Brain Region | Baseline | UoI | Reduction Factor | $p$-value | $d$ |
| --- | --- | --- | --- | --- | --- |
| AC | $0.30 \pm 0.07$ | $0.13 \pm 0.03$ | 2.24 | $5 \times 10^{-23}$ | 1.74 |
| VC 1 | $0.59 \pm 0.11$ | $0.27 \pm 0.07$ | 2.21 | $2 \times 10^{-19}$ | 2.26 |
| VC 2 | $0.66 \pm 0.17$ | $0.26 \pm 0.11$ | 2.56 | $2 \times 10^{-16}$ | 1.87 |
| VC 3 | $0.59 \pm 0.10$ | $0.28 \pm 0.07$ | 2.13 | $2 \times 10^{-20}$ | 2.57 |
| MC 1 | $0.43 \pm 0.12$ | $0.07 \pm 0.04$ | 5.85 | $3 \times 10^{-24}$ | 2.49 |
| MC 2 | $0.46 \pm 0.12$ | $0.08 \pm 0.04$ | 5.50 | $3 \times 10^{-25}$ | 2.37 |
| MC 3 | $0.46 \pm 0.10$ | $0.09 \pm 0.04$ | 5.15 | $1 \times 10^{-25}$ | 2.58 |

**Table B.1c:** Predictive performances

| Brain Region | Baseline | UoI | $p$-value | $d$ |
| --- | --- | --- | --- | --- |
| AC | $0.98 \pm 0.02$ | $0.98 \pm 0.02$ | $5 \times 10^{-4}$ | 0.005 |
| VC 1 | $203 \pm 35$ | $205 \pm 36$ | $3 \times 10^{-16}$ | 0.05 |
| VC 2 | $166 \pm 44$ | $168 \pm 43$ | $3 \times 10^{-12}$ | 0.03 |
| VC 3 | $213 \pm 42$ | $217 \pm 41$ | $2 \times 10^{-18}$ | 0.05 |
| MC 1 | $232 \pm 65$ | $233 \pm 65$ | $1 \times 10^{-23}$ | 0.03 |
| MC 2 | $263 \pm 60$ | $266 \pm 59$ | $1 \times 10^{-23}$ | 0.03 |
| MC 3 | $248 \pm 57$ | $250 \pm 58$ | $2 \times 10^{-25}$ | 0.03 |

**Table B.1d:** Bayesian information criterion

| Brain Region | Baseline | UoI | Median Difference |
| --- | --- | --- | --- |
| AC | $-6240 \pm 1884$ | $-6491 \pm 1859$ | 170 |
| VC 1 | $-15349 \pm 20946$ | $-15516 \pm 20970$ | 149 |
| VC 2 | $-2864 \pm 11046$ | $-3007 \pm 11087$ | 131 |
| VC 3 | $-35164 \pm 34069$ | $-35309 \pm 34066$ | 161 |
| MC 1 | $481 \pm 124$ | $321 \pm 59$ | 162 |
| MC 2 | $542 \pm 150$ | $354 \pm 75$ | 186 |
| MC 3 | $564 \pm 120$ | $362 \pm 67$ | 190 |

### B.2 Encoding model dataset summary and fitted model statistics

Dataset summary (Table B.2a) provides details on the datasets used to fit encoding models, including the number of units and samples across dataset. The following tables provide statistics summarizing aspects of the fitted baseline and UoI models, including selection ratio (Table B.2b), predictive performance (Table B.2c), and Bayesian information criterion (Table B.2d).

**Table B.2a:** Dataset summary

| Brain Region | Number of Units | Number of Samples |
| --- | --- | --- |
| Retina | 125 | 4200 |
| AC | 23 | 89896 |

**Table B.2b:**
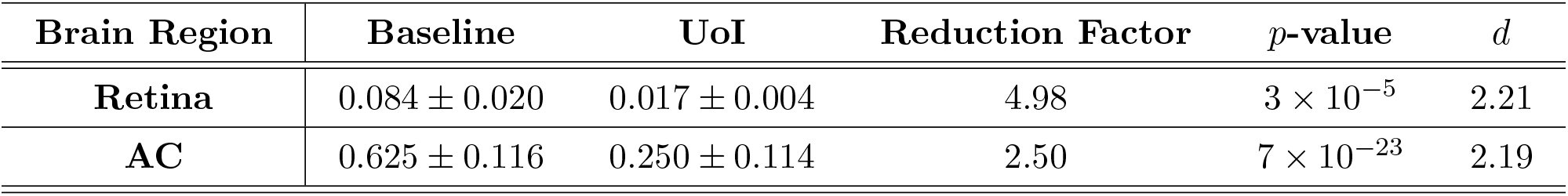
Selection ratio

| Brain Region | Baseline | UoI | Reduction Factor | $p$-value | $d$ |
| --- | --- | --- | --- | --- | --- |
| Retina | $0.084 \pm 0.020$ | $0.017 \pm 0.004$ | 4.98 | $3 \times 10^{-5}$ | 2.21 |
| AC | $0.625 \pm 0.116$ | $0.250 \pm 0.114$ | 2.50 | $7 \times 10^{-23}$ | 2.19 |

**Table B.2c:**
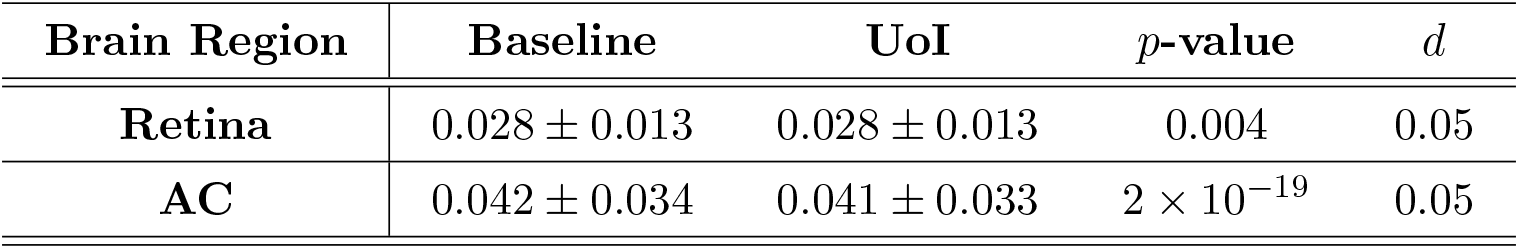
Predictive performances

**Table B.2d:** Bayesian information criterion

| Brain Region | Baseline | UoI | Median Difference |
| --- | --- | --- | --- |
| Retina | $-1646606 \pm 44936$ | $-1647261 \pm 44857$ | 654 |
| AC | $4371 \pm 2495$ | $4351 \pm 2493$ | 19.4 |

### B.3 Decoding model dataset summary and fitted model statistics

Dataset summary (Table B.3a) provides details on the datasets used to fit decoding models, including the number of units and samples across dataset. The following tables provide statistics summarizing aspects of the fitted baseline and UoI models, including selection ratio (Table B.3b), and predictive performance (Table B.3c).

**Table B.3a:** Dataset summary

| Brain Region | Number of Units | Number of Samples |
| --- | --- | --- |
| GPe | 18 | 186 |
| SNr | 36 | 186 |

**Table B.3b:** Selection ratio

| Brain Region | Baseline | UoI | Reduction Factor |
| --- | --- | --- | --- |
| GPe | $0.722 \pm 0.083$ | $0.333 \pm 0.028$ | 2.167 |
| SNr | $0.222 \pm 0.167$ | $0.056 \pm 0.014$ | 4 |

**Table B.3c:** Predictive performances

| Brain Region | Baseline | UoI |
| --- | --- | --- |
| GPe | $0.676 \pm 0.054$ | $0.676 \pm 0.031$ |
| SNr | $1.000 \pm 0.014$ | $1.000 \pm 0.014$ |

## C Frequency response area analysis for tuned and non-tuned electrodes

**Figure C:**
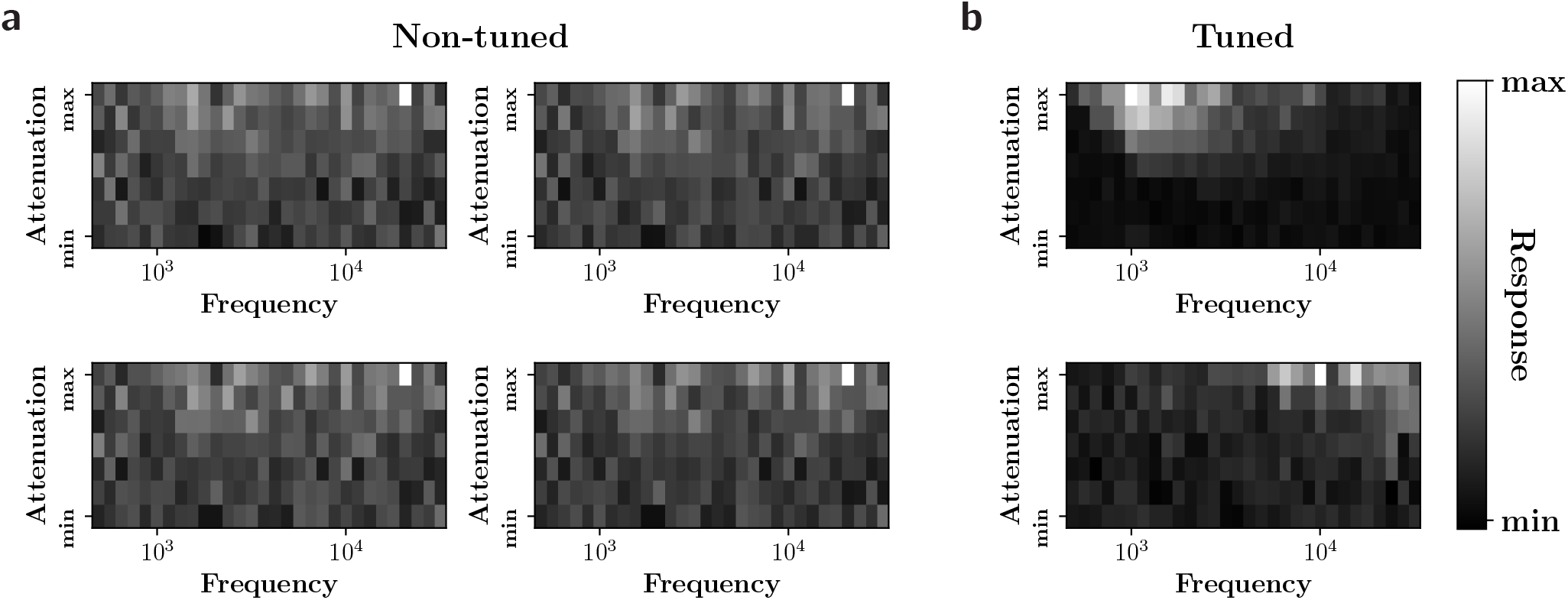
Frequency response area (FRA) analysis of non-tuned electrodes, as determined by UoI, confirm that a frequency tuning model captures no discernible structure in their responses. Each plot depicts the FRA, or the mean responses, across trials, to frequency-attenuation stimulus pairings. The plots are normalized to their maximum value. In each plot, the *y*-axis denote attenuation (ranging from −70 db to 0 db) while the *x*-axis denotes frequency (ranging from 500 Hz to 32 kHz). **a.** FRAs corresponding to the four electrodes that UoI determined to be non-tuned (pink points in Fig 6). **b.** FRAs for two randomly chosen tuned electrodes. The non-tuned electrodes exhibit no discernible structure in the FRAs (in contrast to the clear structure depicted by the tuned electrodes), validating that UoI correctly determined them to be non-tuned.

## References

1. Sejnowski, T. J., Churchland, P. S. & Movshon, J. A. Putting big data to good use in neuroscience. Nature neuroscience 17, 1440 (2014).

2. Marx, V. Biology: The big challenges of big data 2013.

3. Stevenson, I. H. & Kording, K. P. How advances in neural recording affect data analysis. Nature neuroscience 14, 139 (2011).

4. Bzdok, D. & Yeo, B. T. Inference in the age of big data: Future perspectives on neuroscience. Neuroimage 155, 549–564 (2017).

5. Stevenson, I. H., Rebesco, J. M., Miller, L. E. & Körding, K. P. Inferring functional connections between neurons. Current opinion in neurobiology 18, 582–588 (2008).

6. Okatan, M., Wilson, M. A. & Brown, E. N. Analyzing functional connectivity using a network likelihood model of ensemble neural spiking activity. Neural computation 17, 1927–1961 (2005).

7. Truccolo, W., Eden, U. T., Fellows, M. R., Donoghue, J. P. & Brown, E. N. A point process framework for relating neural spiking activity to spiking history, neural ensemble, and extrinsic covariate effects. Journal of neurophysiology 93, 1074–1089 (2005).

8. Pillow, J. W. et al. Spatio-temporal correlations and visual signalling in a complete neuronal population. Nature 454, 995 (2008).

9. Paninski, L., Pillow, J. & Lewi, J. in Computational Neuroscience: Theoretical Insights into Brain Function 493–507 (Elsevier, 2007).

10. Kass, R. E. et al. Computational Neuroscience: Mathematical and Statistical Perspectives. Annual Review of Statistics and Its Application 5, 183–214 (2018).

11. Babadi, B., Casti, A., Xiao, Y., Kaplan, E. & Paninski, L. A generalized linear model of the impact of direct and indirect inputs to the lateral geniculate nucleus. Journal of Vision 10, 22–22 (2010).

12. Zhao, M. et al. An L 1-regularized logistic model for detecting short-term neuronal interactions. Journal of computational neuroscience 32, 479–497 (2012).

13. Stevenson, I. H. et al. Functional Connectivity and Tuning Curves in Populations of Simultaneously Recorded Neurons. PLoS Computational Biology 8 (2012).

14. Song, D. et al. Identification of sparse neural functional connectivity using penalized likelihood estimation and basis functions. Journal of computational neuroscience 35, 335–357 (2013).

15. Bassett, D. S. & Sporns, O. Network neuroscience. Nature neuroscience 20, 353 (2017).

16. Bullmore, E. & Sporns, O. Complex brain networks: graph theoretical analysis of structural and functional systems. Nature reviews neuroscience 10, 186–198 (2009).

17. Bassett, D. S. & Bullmore, E. Small-world brain networks. The neuroscientist 12, 512–523 (2006).

18. Barab±si, A.-L. & Albert, R. Emergence of scaling in random networks. science 286, 509–512 (1999).

19. Melozzi, F. et al. Individual structural features constrain the mouse functional connectome. Proceedings of the National Academy of Sciences 116, 26961–26969 (2019).

20. Friston, K., Moran, R. & Seth, A. K. Analysing connectivity with Granger causality and dynamic causal modelling. Current opinion in neurobiology 23, 172–178 (2013).

21. Seth, A. K., Barrett, A. B. & Barnett, L. Granger causality analysis in neuroscience and neuroimaging. Journal of Neuroscience 35, 3293–3297 (2015).

22. Kiani, R., Cueva, C. J., Reppas, J. B. & Newsome, W. T. Dynamics of neural population responses in prefrontal cortex indicate changes of mind on single trials. Current Biology 24, 1542–1547 (2014).

23. Fulcher, B. D. & Fornito, A. A transcriptional signature of hub connectivity in the mouse connectome. Proceedings of the National Academy of Sciences 113, 1435–1440 (2016).

24. Arnemann, K. L., Stöber, F., Narayan, S., Rabinovici, G. D. & Jagust, W. J. Metabolic brain networks in aging and preclinical Alzheimer’s disease. NeuroImage: Clinical 17, 987–999 (2018).

25. Dayan, P. & Abbott, L. F. Theoretical neuroscience: computational and mathematical modeling of neural systems (MIT press, 2001).

26. Schwartz, O., Pillow, J. W., Rust, N. C. & Simoncelli, E. P. Spike-triggered neural characterization. Journal of vision 6, 13–13 (2006).

27. Triplett, M. A. & Goodhill, G. J. Probabilistic Encoding Models for Multivariate Neural Data. Frontiers in Neural Circuits 13, 1 (2019).

28. Sharpee, T. O. Computational Identification of Receptive Fields. Annual Review of Neuroscience 36, 103–120 (2013).

29. Theunissen, F. et al. Estimating spatio-temporal receptive fields of auditory and visual neurons from their responses to natural stimuli. Network: Computation in Neural Systems 12, 289–316 (2001).

30. Hubel, D. H. & Wiesel, T. N. Receptive fields, binocular interaction and functional architecture in the cat’s visual cortex. The Journal of Physiology 160, 106–154 (1962).

31. Rao, R. P. & Ballard, D. H. Predictive coding in the visual cortex: a functional interpretation of some extra-classical receptive-field effects. Nature neuroscience 2, 79–87 (1999).

32. Vinje, W. E. & Gallant, J. L. Sparse coding and decorrelation in primary visual cortex during natural vision. Science 287, 1273–1276 (2000).

33. Zhu, M. & Rozell, C. J. Visual nonclassical receptive field effects emerge from sparse coding in a dynamical system. PLoS computational biology 9 (2013).

34. Glaser, J. I., Chowdhury, R. H., Perich, M. G., Miller, L. E. & Kording, K. P. Machine learning for neural decoding. arXiv preprint arXiv:1708.00909 (2017).

35. Naselaris, T., Kay, K. N., Nishimoto, S. & Gallant, J. L. Encoding and decoding in fMRI. Neuroimage 56, 400–410 (2011).

36. Holdgraf, C. R. et al. Encoding and decoding models in cognitive electrophysiology. Frontiers in systems neuroscience 11, 61 (2017).

37. Bouchard, K. E., Mesgarani, N., Johnson, K. & Chang, E. F. Functional organization of human sensorimotor cortex for speech articulation. Nature 495, 327 (2013).

38. Bouchard, K. E. & Chang, E. F. Neural decoding of spoken vowels from human sensory-motor cortex with high-density electrocorticography in 2014 36th Annual International Conference of the IEEE Engineering in Medicine and Biology Society (2014), 6782–6785.

39. Livezey, J. A., Bouchard, K. E. & Chang, E. F. Deep learning as a tool for neural data analysis: speech classification and cross-frequency coupling in human sensorimotor cortex. PLoS computational biology 15, e1007091 (2019).

40. Anumanchipalli, G. K., Chartier, J. & Chang, E. F. Speech synthesis from neural decoding of spoken sentences. Nature 568, 493–498 (2019).

41. Yamashita, O., Sato, M.-a., Yoshioka, T., Tong, F. & Kamitani, Y. Sparse estimation automatically selects voxels relevant for the decoding of fMRI activity patterns. NeuroImage 42, 1414–1429 (2008).

42. Rich, E. L. & Wallis, J. D. Decoding subjective decisions from orbitofrontal cortex. Nature Neuroscience 19, 973–980. https://doi.org/10.1038/nn.4320 (2016).

43. Pandarinath, C. et al. Inferring single-trial neural population dynamics using sequential auto-encoders. Nature methods 15, 805–815 (2018).

44. Wander, J. D. et al. Distributed cortical adaptation during learning of a brain–computer interface task. Proceedings of the National Academy of Sciences 110, 10818–10823 (2013).

45. Carmena, J. M. et al. Learning to control a brain–machine interface for reaching and grasping by primates. PLoS biology 1 (2003).

46. Schwarz, G. Estimating the Dimension of a Model. Ann. Statist. 6, 461–464 (Mar. 1978).

47. Tibshirani, R. Regression shrinkage and selection via the lasso. Journal of the Royal Statistical Society: Series B (Methodological) 58, 267–288 (1996).

48. Hastie, T., Tibshirani, R. & Wainwright, M. Statistical learning with sparsity: the lasso and generalizations (Chapman and Hall/CRC, 2015).

49. Yu, B. et al. Stability. Bernoulli 19, 1484–1500 (2013).

50. Bousquet, O. & Elisseeff, A. Stability and generalization. Journal of machine learning research 2, 499–526 (2002).

51. Lim, C. & Yu, B. Estimation stability with cross-validation (ESCV). Journal of Computational and Graphical Statistics 25, 464–492 (2016).

52. Baldassarre, L., Pontil, M. & Mourão-Miranda, J. Sparsity is better with stability: Combining accuracy and stability for model selection in brain decoding. Frontiers in neuroscience 11, 62 (2017).

53. Bouchard, K. et al. in Advances in Neural Information Processing Systems 30 (eds Guyon, I. et al.) 1078–1086 (Curran Associates, Inc., 2017).

54. Ubaru, S., Wu, K. & Bouchard, K. E. UoI-NMF Cluster: A Robust Nonnegative Matrix Factorization Algorithm for Improved Parts-Based Decomposition and Reconstruction of Noisy Data in 2017 16th IEEE International Conference on Machine Learning and Applications (ICMLA) (Dec. 2017), 241–248.

55. Sachdeva, P., Livezey, J., Tritt, A. & Bouchard, K. PyUoI: The Union of Intersections Framework in Python. Journal of Open Source Software 4, 1799 (2019).

56. Friedman, J., Hastie, T. & Tibshirani, R. Regularization paths for generalized linear models via coordinate descent. Journal of statistical software 33, 1 (2010).

57. Breiman, L. Bagging predictors. Machine learning 24, 123–140 (1996).

58. Friedman, J. H. Greedy function approximation: a gradient boosting machine. Annals of statistics, 1189–1232 (2001).

59. Nelder, J. A. & Wedderburn, R. W. Generalized linear models. Journal of the Royal Statistical Society: Series A (General) 135, 370–384 (1972).

60. Fan, J. & Li, R. Variable selection via nonconcave penalized likelihood and its oracle properties. Journal of the American statistical Association 96, 1348–1360 (2001).

61. Bouchard, K. E. Bootstrapped adaptive threshold selection for statistical model selection and estimation. arXiv preprint arXiv:1505.03511 (2015).

62. Javanmard, A. & Montanari, A. Confidence intervals and hypothesis testing for high-dimensional regression. The Journal of Machine Learning Research 15, 2869–2909 (2014).

63. Dougherty, M. E., Nguyen, A. P. Q., Baratham, V. L. & Bouchard, K. E. Laminar origin of evoked ECoG high-gamma activity in 2019 41st Annual International Conference of the IEEE Engineering in Medicine and Biology Society (EMBC) (July 2019), 4391–4394.

64. Teeters, J. L., Harris, K. D., Millman, K. J., Olshausen, B. A. & Sommer, F. T. Data Sharing for Computational Neuroscience. Neuroinformatics 6, 47–55 (Mar. 2008).

65. Kohn, A. & Smith, M. A. Utah array extracellular recordings of spontaneous and visually evoked activity from anesthetized macaque primary visual cortex (V1) 2016. http://dx.doi.org/10.6080/K0NC5Z4X.

66. Smith, M. A. & Kohn, A. Spatial and Temporal Scales of Neuronal Correlation in Primary Visual Cortex. Journal of Neuroscience 28, 12591–12603 (2008).

67. Kelly, R. C., Smith, M. A., Kass, R. E. & Lee, T. S. Local field potentials indicate network state and account for neuronal response variability. Journal of computational neuroscience 29, 567–579 (2010).

68. O’Doherty, J. E., Cardoso, M. M. B., Makin, J. G. & Sabes, P. N. Nonhuman Primate Reaching with Multichannel Sensorimotor Cortex Electrophysiology May 2017. https://doi.org/10.5281/zenodo.583331.

69. Makin, J. G., O’Doherty, J. E., Cardoso, M. M. B. & Sabes, P. N. Superior arm-movement decoding from cortex with a new, unsupervised-learning algorithm. Journal of Neural Engineering 15, 026010 (Jan. 2018).

70. Zhang, Y.-F., Asari, H. & Meister, M. Multi-electrode recordings from retinal ganglion cells 2014. http://dx.doi.org/10.6080/K0RF5RZT.

71. Lefebvre, J. L., Zhang, Y., Meister, M., Wang, X. & Sanes, J. R. *γ*-Protocadherins regulate neuronal survival but are dispensable for circuit formation in retina. Development 135, 4141–4151 (2008).

72. Gu, B.-M., Schmidt, R. & Berke, J. D. Globus pallidus dynamics reveal covert strategies for behavioral inhibition. bioRxiv (2020).

73. Pedregosa, F. et al. Scikit-learn: Machine Learning in Python. Journal of Machine Learning Research 12, 2825–2830 (2011).

74. Buitinck, L. et al. API design for machine learning software: experiences from the scikit-learn project in ECML PKDD Workshop: Languages for Data Mining and Machine Learning (2013), 108–122.

75. Shao, J. Linear model selection by cross-validation. Journal of the American statistical Association 88, 486–494 (1993).

76. Shao, J. An asymptotic theory for linear model selection. Statistica sinica, 221–242 (1997).

77. Zhang, Y., Li, R. & Tsai, C.-L. Regularization parameter selections via generalized information criterion. Journal of the American Statistical Association 105, 312–323 (2010).

78. Gong, P. & Ye, J. A Modified Orthant-Wise Limited Memory Quasi-Newton Method with Convergence Analysis in Proceedings of the 32nd International Conference on International Conference on Machine Learning-Volume 37 (2015), 276–284.

79. Wilcoxon, F. in Breakthroughs in statistics 196–202 (Springer, 1992).

80. Neath, A. A. & Cavanaugh, J. E. The Bayesian information criterion: background, derivation, and applications. Wiley Interdisciplinary Reviews: Computational Statistics 4, 199–203 (2012).

81. Cohen, J. Statistical power analysis for the behavioral sciences (Academic press, 2013).

82. Sawilowsky, S. S. New effect size rules of thumb. Journal of Modern Applied Statistical Methods 8, 26 (2009).

83. Wang, H. E. et al. A systematic framework for functional connectivity measures. Frontiers in neuroscience 8, 405 (2014).

84. Newman, M. E. Modularity and community structure in networks. Proceedings of the national academy of sciences 103, 8577–8582 (2006).

85. Clauset, A., Newman, M. E. & Moore, C. Finding community structure in very large networks. Physical review E 70, 066111 (2004).

86. Watts, D. J. & Strogatz, S. H. Collective dynamics of ‘small-world’networks. nature 393, 440 (1998).

87. Bassett, D. S. & Bullmore, E. T. Small-world brain networks revisited. The Neuroscientist 23, 499–516 (2017).

88. Telesford, Q. K., Joyce, K. E., Hayasaka, S., Burdette, J. H. & Laurienti, P. J. The ubiquity of small-world networks. Brain connectivity 1, 367–375 (2011).

89. Karklin, Y. & Simoncelli, E. P. Efficient coding of natural images with a population of noisy linear-nonlinear neurons in Advances in neural information processing systems (2011), 999–1007.

90. Olshausen, B. A. & Field, D. J. Emergence of simple-cell receptive field properties by learning a sparse code for natural images. Nature 381, 607–609 (1996).

91. Honey, C. J., Kötter, R., Breakspear, M. & Sporns, O. Network structure of cerebral cortex shapes functional connectivity on multiple time scales. Proceedings of the National Academy of Sciences 104, 10240–10245 (2007).

92. Markov, N. T. et al. Cortical high-density counterstream architectures. Science 342, 1238406 (2013).

93. Das, A. & Fiete, I. R. Systematic errors in connectivity inferred from activity in strongly coupled recurrent circuits. bioRxiv (2019).

94. Song, S., Sjöström, P. J., Reigl, M., Nelson, S. & Chklovskii, D. B. Highly nonrandom features of synaptic connectivity in local cortical circuits. PLoS biology 3 (2005).

95. Barbour, B., Brunel, N., Hakim, V. & Nadal, J.-P. What can we learn from synaptic weight distributions? TRENDS in Neurosciences 30, 622–629 (2007).

96. Balasubramanian, M. et al. Optimizing the Union of Intersections LASSO (UoI-LASSO) and Vector Autoregressive (UoI-VAR) Algorithms for Improved Statistical Estimation at Scale. arXiv:1808.06992 (2018).

97. Ruiz, T., Balasubramanian, M., Bouchard, K. E. & Bhattacharyya, S. Sparse, Low-bias, and Scalable Estimation of High Dimensional Vector Autoregressive Models via Union of Intersections. arXiv:1908.11464 (2019).

98. Gu, S. et al. Controllability of structural brain networks. Nature communications 6, 1–10 (2015).

99. Stringer, C., Pachitariu, M., Steinmetz, N., Carandini, M. & Harris, K. D. High-dimensional geometry of population responses in visual cortex. Nature 571, 361–365 (2019).

100. Schüz, A., Chaimow, D., Liewald, D. & Dortenman, M. Quantitative aspects of corticocortical connections: a tracer study in the mouse. Cerebral cortex 16, 1474–1486 (2006).

101. Isely, G., Hillar, C. & Sommer, F. Deciphering subsampled data: adaptive compressive sampling as a principle of brain communication in Advances in neural information processing systems (2010), 910–918.

102. Kohn, A., Coen-Cagli, R., Kanitscheider, I. & Pouget, A. Correlations and neuronal population information. Annual review of neuroscience 39, 237–256 (2016).

103. Wang, H., Li, R. & Tsai, C.-L. Tuning parameter selectors for the smoothly clipped absolute deviation method. Biometrika 94, 553–568 (2007).

104. Akaike, H. A new look at the statistical model identification. IEEE transactions on automatic control 19, 716–723 (1974).

105. Rissanen, J. Modeling by shortest data description. Automatica 14, 465–471 (1978).

106. George, E. & Foster, D. P. Calibration and empirical Bayes variable selection. Biometrika 87, 731–747 (2000).

107. Vidne, M. et al. Modeling the impact of common noise inputs on the network activity of retinal ganglion cells. Journal of computational neuroscience 33, 97–121 (2012).

108. Okun, M. et al. Diverse coupling of neurons to populations in sensory cortex. Nature 521, 511–515 (2015).

109. Byron, M. Y. et al. Gaussian-process factor analysis for low-dimensional single-trial analysis of neural population activity in Advances in neural information processing systems (2009), 1881–1888.

110. Paninski, L. et al. A new look at state-space models for neural data. Journal of computational neuroscience 29, 107–126 (2010).

111. Macke, J. H. et al. Empirical models of spiking in neural populations in Advances in neural information processing systems (2011), 1350–1358.

112. Cunningham, J. P. & Byron, M. Y. Dimensionality reduction for large-scale neural recordings. Nature neuroscience 17, 1500 (2014).

113. Kell, A. J. & McDermott, J. H. Deep neural network models of sensory systems: windows onto the role of task constraints. Current opinion in neurobiology 55, 121–132 (2019).

114. Abbasi-Asl, R. & Yu, B. Structural Compression of Convolutional Neural Networks Based on Greedy Filter Pruning 2017. arXiv: 1705.07356.

115. Zhang, C.-H. et al. Nearly unbiased variable selection under minimax concave penalty. The Annals of statistics 38, 894–942 (2010).

116. Kitano, H. Computational systems biology. Nature 420, 206–210 (2002).

117. Basu, S., Kumbier, K., Brown, J. B. & Yu, B. Iterative random forests to discover predictive and stable high-order interactions. Proceedings of the National Academy of Sciences 115, 1943–1948 (2018).

118. Jaqaman, K. & Danuser, G. Linking data to models: data regression. Nature Reviews Molecular Cell Biology 7, 813–819 (2006).

119. Suárez, E., Párez, C. M., Rivera, R. & Martinez, M. N. Applications of Regression Models in Epidemiology (John Wiley & Sons, 2017).

120. Guisan, A., Edwards Jr, T. C. & Hastie, T. Generalized linear and generalized additive models in studies of species distributions: setting the scene. Ecological modelling 157, 89–100 (2002).

